# Cariprazine modulates intrinsic excitability and network dynamics of hippocampal neurons in a cell-type dependent manner

**DOI:** 10.64898/2026.05.22.727184

**Authors:** Melinda E. Gazdik, Iván Fejes, Ádám Tiszlavicz, Anna A. Abbas, Lea Danics, Balázs Kis, Anna Ország, Kai Kummer, Michaela Kress, Katalin Schlett, János M. Réthelyi, András Benczúr, Karri P. Lamsa, Attila Szűcs, Karolina Pircs

**Affiliations:** Institute of Clinical Pathophysiology, Semmelweis University, 1094 Budapest, Hungary; Hungarian Centre of Excellence for Molecular Medicine - Semmelweis University (HCEMM-SU), Neurobiology and Neurodegenerative Diseases Research Group, 1094 Budapest, Hungary; HUN-REN-SZTAKI-SU Rejuvenation Research Group, HUN-REN Office for Supported Research Groups (TKI), 1052 Budapest, Hungary; Department of Physiology and Neurobiology, Institute of Biology, Eötvös Loránd University, 1117 Budapest, Hungary; Institute for Computer Science and Control (SZTAKI), Hungarian Research Network (HUN-REN), 1111 Budapest, Hungary; Budapest University of Technology and Economics, 1111 Budapest, Hungary; Hungarian Centre of Excellence for Molecular Medicine (HCEMM), Human Neuron Physiology and Therapy Core Group, 6728 Szeged, Hungary; Institute of Physiology, Medical University of Innsbruck, 6020 Innsbruck, Austria; Department of Psychiatry and Psychotherapy, Semmelweis University, 1083 Budapest, Hungary

**Keywords:** cariprazine, schizophrenia, hippocampus, antipsychotic medications, real-world evidence, multielectrode array, whole-cell patch clamp, electrophysiology, Kv1 channels

## Abstract

Schizophrenia is a severe psychiatric disorder associated with altered dopaminergic signaling and hippocampal circuit dysfunction. Although antipsychotic medications remain the standard treatment, many are limited by incomplete efficacy and adverse effects. Cariprazine, a dopamine D2/D3 receptor partial agonist, has a favorable clinical profile, but its effects on neuronal excitability and network activity remain incompletely understood. Here, we integrated nationwide real-world clinical data with *in vitro* electrophysiology, computational modeling, and molecular analyses to define the neuronal actions of cariprazine. Among Hungarian patients diagnosed with schizophrenia and receiving index-drug monotherapy with one of the three prespecified D2/D3 targeting antipsychotics, haloperidol was associated with worse survival and a higher cumulative incidence of first registered suicide attempt than cariprazine or aripiprazole in matched observational cohorts. In primary mouse hippocampal cultures, multielectrode array recordings showed that acute cariprazine treatment moderately reduced spontaneous firing in a dose-dependent manner and prolonged burst intervals while largely preserving network synchronization. These effects were milder than those of haloperidol and aripiprazole. Whole-cell patch-clamp recordings revealed cell-type-dependent effects, with reduced intrinsic excitability and increased firing irregularity mainly in regular- and stuttering-type neurons. Conductance-based modeling identified enhanced Kv1-mediated D-type potassium currents as sufficient to reproduce these effects. Consistent with this mechanism, chronic cariprazine treatment altered Kv1.2 protein distribution without changing *Kcna2*/*Kcna3* or *Drd1*/*Drd2*/*Drd3* transcript expression. These findings identify modulation of intrinsic excitability via Kv1/D-type potassium currents as a candidate cellular mechanism of cariprazine and provide a translational link between real-world evidence and circuit-level drug effects.

## INTRODUCTION

Neuropsychiatric disorders represent a major global health burden and are among the leading causes of disability worldwide, with an increasing trend over the past three decades (1, 2). These conditions are highly heterogeneous in etiology, symptomatology, disease course, and frequently involve cognitive impairment and affective dysregulation (3). Schizophrenia is one of the most severe psychiatric disorders, typically emerging in early adulthood and characterized by a chronic, relapsing course that profoundly impairs social and occupational functioning. Clinically, schizophrenia is defined by three core symptom domains: positive symptoms (including hallucinations, delusions, disorganized behavior), negative symptoms (such as social withdrawal, anhedonia, and alogia), and cognitive symptoms encompassing deficits in attention, working memory, and executive function (4). While positive symptoms are often responsive to pharmacotherapy, negative and cognitive symptoms, which are strong predictors of long-term functional impairment remain largely untreatable in terms of pharmacological interventions (4, 5).

Current pharmacological treatments for schizophrenia primarily target dopaminergic signaling, particularly dopamine D2 receptors (6). First-generation, typical antipsychotics, such as haloperidol, are effective against positive symptoms but are associated with substantial extrapyramidal side effects and limited efficacy for negative and cognitive symptom domains (7). Second- and third-generation, atypical antipsychotics were developed to improve tolerability and broaden therapeutic efficacy by involving serotonin 2A (5-HT_2A_) receptor antagonism; however, many still fail to meaningfully address negative symptoms and cognitive impairment, and their mechanisms of action at the level of neuronal circuits and intrinsic cellular properties are not fully understood (6–8). There is a critical need for a better understanding of how newer antipsychotic compounds modulate neuronal function beyond canonical receptor-level pharmacology, particularly within brain regions supporting cognition and network coordination.

Cariprazine is a third-generation antipsychotic approved for the treatment of schizophrenia, bipolar affective disorder, and major depressive disorder (9). Pharmacologically, cariprazine is distinguished by its high affinity for dopamine D3 receptors and partial agonist activity at both D2 and D3 receptors, exhibiting a greater preference for D3 receptors than other partial D2/D3 agonists such as aripiprazole and brexpiprazole. In line with these clinical observations, positron emission tomography (PET) studies demonstrate a marked *in vivo* preference of cariprazine for dopamine D3 over D2 receptors, distinguishing it pharmacologically from other partial agonists (10).

In addition, cariprazine displays partial agonism at serotonin 5-HT1A receptors, low-to-moderate affinity for 5-HT2A, 5-HT2C, and histamine H1 receptors, and potent antagonism at 5-HT2B receptors (11). Clinically, cariprazine has demonstrated efficacy across positive and negative symptom domains with a comparatively favorable side-effect profile; however, the cellular and network-level mechanisms underlying these effects remain poorly characterized.

Converging evidence from real-world clinical data and meta-analyses further highlights the need to investigate whether cariprazine engages neuronal mechanisms that are distinct from those of other antipsychotic agents. Based on real-world evidence, second-generation antipsychotics are associated with significantly lower all-cause mortality compared with first-generation compounds, as demonstrated in large meta-analyses encompassing more than 300,000 patients (12). Although mortality outcomes have not yet been evaluated specifically for cariprazine, real-world observational studies indicate that cariprazine treatment is associated with favorable healthcare utilization profiles, including fewer all-cause and mental health-related hospitalizations compared with other atypical antipsychotics (13, 14). In addition, observational and randomized evidence suggests that cariprazine provides particular benefit for negative symptom domains - such as avolition, anhedonia, and blunted affect - which are closely linked to motivational and reward-processing circuits and remain inadequately treated by most available agents (15, 16).

At the circuit level, the hippocampus is a critical integrative hub for memory, cognitive coordination, and emotional and motivational regulation, functions that are consistently disrupted in schizophrenia. Both clinical and preclinical evidence implicates hippocampal network dysfunction in the disorder, including altered excitation-inhibition balance, disrupted gamma-band oscillations, and abnormalities in GABAergic interneuron function (17–19). While antipsychotic drugs are known to influence hippocampal activity, their effects on network dynamics and intrinsic neuronal excitability remain undefined. Notably, recent preclinical studies suggest that cariprazine can normalize pathological hippocampal gamma oscillations and modulate neurotransmitter signaling in a D3 receptor-dependent manner, indicating activity- and state-dependent circuit stabilization rather than uniform suppression (20, 21). However, the cellular substrates linking these network effects to intrinsic excitability and cell-type-specific mechanisms remain largely unexplored.

In the present study, we first identified patients with schizophrenia from nationwide real-world data and then compared outcomes from 1^st^ January 2019 to 31^st^ December 2020 among patients receiving haloperidol, cariprazine, or aripiprazole index-drug monotherapy. This analysis demonstrated a clear distinction between patients treated with haloperidol and those receiving the two partial agonists, with cariprazine and aripiprazole exhibiting more favorable clinical outcome profiles. We then used primary hippocampal cultures prepared from CD-1 mice to investigate the neuronal mechanisms that may contribute to these differences, combining acute and prolonged drug exposure paradigms with network-level electrophysiology, intrinsic cellular recordings, computational modeling, and molecular validation. By integrating real-world clinical observations with circuit-, cellular-, and molecular-level analyses, our work aims to provide mechanistic insight into how cariprazine modulates hippocampal neuronal function.

Together, this study links population-level clinical observations with experimental neurobiology to advance the mechanistic understanding of third-generation antipsychotics. Our findings identify noncanonical, cell-type- and ion channel-associated effects of cariprazine that extend beyond classical dopaminergic signaling. These findings provide a translational framework for understanding how modulation of hippocampal circuitry may contribute to improved clinical outcomes in schizophrenia.

## MATERIALS AND METHODS

### Real-world clinical data

#### Real-world data source and study design

To compare clinical outcomes associated with antipsychotic agents acting primarily on dopamine D2/D3 receptors, we conducted a retrospective, nationwide, register-based, comparative cohort study using routinely collected administrative healthcare data from the Hungarian National Health Insurance Fund Administration (NHIFA; Hungarian acronym: NEAK). The NHIFA database captures all publicly financed healthcare services in Hungary between 2010 and 2021 and links de-identified individual-level information from five major sources; primary care, outpatient specialty care, hospital utilization, a demographic registry based on social security numbers, and dispensed prescription records. Diagnoses were recorded using the 10^th^ Revision of the International Classification of Diseases (ICD-10), and medication dispensing records were classified according to Anatomical Therapeutic Chemical (ATC) codes. The demographic registry included date of birth, sex, date of death, citizenship, and social security number validity. Personal identifiers were converted into depersonalized but linkable identifiers by National Infocommunications Ltd, allowing longitudinal follow-up while preserving patient anonymity. Data access was restricted to a secure research environment without internet connectivity, protected by strict physical access controls.

#### Study cohort

The study cohort comprised patients with schizophrenia-spectrum disorders identified in the NHIFA database who had at least one primary ICD-10 diagnosis within the F2.x category recorded during the index period from 1^st^ January 2018 to 31^st^ December 2018. Eligible patients were required to have received at least one dispensed prescription for one of the study drugs of interest (haloperidol, aripiprazole, or cariprazine) during the same period. To ensure mutually exclusive exposure groups, only patients who received prescriptions for a single study drug of interest were included, although patients could also have received other antipsychotic medications during the index period. To reduce diagnostic heterogeneity and confounding related to substance use, patients with any recorded ICD-10 F1.x diagnosis during the index period were excluded. Patients were also excluded if pregnancy-related diagnostic codes were recorded during the index period. In addition, only patients who were alive at the end of the index period were eligible for inclusion. Numerical details of the cohort are provided in Supplementary Table 1 and Supplementary Table S6.

#### Exposure definition

Antipsychotic exposure was defined on the basis of dispensed prescriptions identified using ATC codes during the index period. The study drugs of interest were haloperidol (first-generation antipsychotic; dopamine D2 receptor antagonist; ATC code N05AD01), aripiprazole (dopamine D2/D3 partial agonist; ATC code N05AX12), and cariprazine (third-generation antipsychotic; D3-preferring dopamine D2/D3 partial agonist; ATC code N05AX15). Patients were assigned to exposure groups according to the study drug of interest dispensed during the index period. As cariprazine was authorized in the European Union in July 2017 and first appeared in our database only in late 2017, the index period was defined to begin on 1^st^ January 2018 rather than at the time of its first observed prescription. This approach allowed time for its integration into routine clinical practice, reduced potential bias arising from early post-launch prescribing patterns, and improved comparability across treatment groups. Throughout the manuscript, we use the term index-drug monotherapy to refer to mutually exclusive exposure to one of the three prespecified index antipsychotics - haloperidol, aripiprazole, or cariprazine - during the index period. This exposure definition was based on the principal study drug used for group assignment; concomitant prescriptions for antipsychotics outside these three index agents were not used to define exposure-group membership and were captured as baseline co-medication covariates in the propensity-score framework.

#### Confounder assessment and covariate balance

To assess and reduce baseline differences between exposure groups, we performed pairwise high-dimensional propensity score matching for each drug comparison. Because the NHIFA database contains only routinely collected administrative healthcare records and related metadata, adjustment was limited to confounders observable within the dataset. We therefore pre-selected clinically relevant comorbidities and co-medications that could influence treatment choice or study outcomes, including major psychiatric and somatic disease groups, as well as medications such as other antipsychotics, benzodiazepines, antidepressants, antiepileptics, antiparkinsonian agents, antidiabetic drugs, lipid-lowering agents, antithrombotic agents, and analgesics. The full list of pre-selected comorbidities and co-medications are provided in the Supplementary Methods.

Pre-defined ICD-10 diagnostic categories were grouped into broader clinically meaningful comorbidity domains, and the occurrence of these grouped diagnoses during the index period was recorded for each patient. Similarly, exposure to the pre-specified ATC drug classes during the index period was captured. These variables formed the core set of covariates for propensity score estimation. To further improve covariate capture, we generated candidate high-dimensional diagnostic and medication features from all 3-character ICD-10 codes and 5-character ATC codes recorded during the index period. Codes were retained as candidate features if they occurred in at least 0.1% of the cohort (corresponding to at least 12 individuals).

Propensity scores were estimated separately for each pairwise exposure comparison using an elastic net model with 5-fold cross-validation. Only pre-specified covariates and the retained high-dimensional diagnostic and medication features were entered into the model. To avoid unstable estimation due to sparse covariates and complete or near-complete separation of the treatment groups, variables were excluded from the model if their prevalence was below 5% in the combined pairwise comparison cohort or below 3% in either treatment group. Patients were then matched based on estimated propensity scores to minimize residual imbalance between treatment groups, with a target standardized mean difference (SMD) of <0.1 for all measured covariates. To maximize the number of matched pairs, we used optimal matching based on the Hungarian algorithm with a composite distance metric incorporating both propensity score differences and Hamming distance between covariate profiles. To further reduce imbalance in key demographic variables, matching was stratified by sex and age, with exact matching on sex and 5-year age bands.

The top 20 features with the highest absolute standardized mean differences for each exposure-group comparison, together with the corresponding propensity score distributions, are shown in Supplementary Fig. 1. The full list of features included in the high-dimensional propensity score models for each exposure-group pair, as well as the standardized mean differences before and after matching, is provided in Supplementary Tables 3-5. After matching, we obtained 2,656 matched pairs in the aripiprazole-haloperidol comparison, 1,119 matched pairs in the cariprazine-haloperidol comparison, and 1,475 matched pairs in the aripiprazole-cariprazine comparison. In each comparison, all features included in the propensity score model had post-matching standardized mean differences below 0.1. Post-matching balance was assessed both for model-included variables and for selected clinically relevant baseline characteristics reported descriptively in Supplementary Table 1, including variables not fully represented in the final matching feature set.

#### Clinical data analysis

All analyses were conducted using time-to-event methods to compare outcomes across exposure groups. The follow-up period was defined from 1^st^ January 2019 through 31^st^ December 2020. The primary outcome was all-cause mortality, defined as death from any cause during follow-up. Secondary outcomes were (i) first psychiatric hospitalization, defined as the first inpatient admission with a schizophrenia-spectrum diagnosis (ICD-10 F2.x), and (ii) first registered suicide attempt, identified by ICD-10 codes X60-X84. For all outcomes, patients who did not experience the event of interest, or any competing event, during follow-up were censored at the end of the follow-up period. Time-to-event was measured in months from the start of follow-up.

For all-cause mortality, Kaplan-Meier curves were estimated and compared between exposure groups using pairwise log-rank tests. For first psychiatric hospitalization and first registered suicide attempt, death was treated as a competing event. Accordingly, cumulative incidence functions were estimated and compared between exposure groups using pairwise Gray’s tests.

#### Software and statistical analysis, notation

All statistical analyses were conducted using Python 3.12. Kaplan-Meier estimation was performed using scikit-survival, whereas log-rank tests and Cox proportional hazards models were fitted using lifelines. Owing to restrictions of the secure local analysis environment, Gray’s test was implemented in Python through a custom reimplementation based on the original Fortran implementation and R wrapper structure of the cmprsk framework. Matching between exposure groups was performed using a custom-developed algorithm based on the Hungarian algorithm for optimal pair matching and a composite distance function incorporating both high-dimensional propensity score differences and the weighted Hamming distance between candidate pairs. To further improve covariate balance, the weighting of individual features in the distance function was optimized using an evolutionary algorithm to minimize the maximum absolute standardized mean difference across covariates. The source code for the matching algorithm and custom statistical functions will be made publicly available on GitHub. Statistical significance was defined as follows: ns, p ≥ 0.05; *, 0.01 ≤ p < 0.05; **, 0.001 ≤ p < 0.01; ***, p < 0.001.

### *In vitro* experimental and computational modeling data

#### Animals

CD-1 wild-type mice (CD-1® IGS Mouse, Ctrl: CD1(ICR); Strain Code: 022; Charles River Laboratories, Wilmington, MA, USA) were housed in a temperature-controlled facility (22 ± 1 °C) under a 12-hour light/dark cycle with ad libitum access to food and water.

Adult male and female mice were paired for breeding and housed together for a 4-day period, during which females were checked daily for the presence of a vaginal plug. Pregnant females were monitored daily throughout gestation until tissue collection.

According to the Hungarian and European Union Animal welfare legislation (in accordance with the Hungarian Act of Animal Care and Experimentation (1998, XXVIII) and with the directive 2010/63/EU of the European Parliament and of the Council of 22 September 2010 on the protection of animals used for scientific purposes), sacrificing animals to obtain brain tissue for primary neuronal cultures is regarded as “organ donation”, therefore, individual ethical approval was not required. All efforts were made to minimize the number of animals used in this study.

#### Cell culture

Primary hippocampal cultures were prepared from mouse embryos obtained from CD-1 wild-type mothers between the embryonic age of (E)16-19, according to (22). Cultures were plated in 24-well CELLSTAR plates (Greiner Bio-One) or 24-well CytoView plates (Axion Biosystems). Plasma-cleaned (Diener Zepto plasma cleaner, 100W, 10 min), rounded glass coverslips (Ø12 mm) were placed into wells designated for whole-cell patch clamp recordings. Prior to plating, wells were surface-coated with poly-L-lysine (PLL; Sigma-Aldrich) and laminin (Sigma-Aldrich) to ensure proper cell adhesion.

Pregnant females were sacrificed, and embryos were used as tissue donors. Brains were removed from the embryos, and hippocampi were dissected and collected. Hippocampal tissue was enzymatically digested using trypsin-EDTA and DNase I in PBS (Gibco, Thermo Fisher Scientific) for 15 minutes at 37 °C. Enzymatic digestion was stopped by addition of heat-inactivated fetal calf serum (FCS), followed by centrifugation at 400 rcf for 10 minutes. The supernatant was discarded, and tissue was mechanically triturated by gentle up-and-down pipetting motions to obtain a single-cell suspension and filtered through a gamma-sterilized polystyrol mesh with 42 µm pore size (EmTek Ltd, Hungary). Viable cells were counted using a Bürker chamber with trypan blue staining, and cells were seeded on the pre-coated surfaces at a density of 70,000 cells/cm^2^ in Neurobasal Plus (Cat. No. A3582901; Thermo Fisher Scientific, Waltham, MA, USA), supplemented with 2% B27 Plus (Cat. No. A3582801; Thermo Fisher Scientific, Waltham, MA, USA), 5% FCS (PAN Biotech, Aidenbach, Germany) and 0.5 mM GlutaMAX (Cat. No. 35050038; Thermo Fisher Scientific, Waltham, MA, USA). On the fifth day after plating (DIV5), one third of the medium was changed to BrainPhys (Cat. No. 05790; StemCell Technologies, Vancouver, BC, Canada) supplemented with 2% NeuroCult SM1 (Cat. No. 05711; StemCell Technologies, Vancouver, BC, Canada).

#### Antipsychotic drug treatments

Cariprazine and other antipsychotic compounds were dissolved in dimethyl sulfoxide (DMSO; CryoMACS, Miltenyi Biotec, Germany), vortexed, and sterile-filtered to generate 10 mM stock solutions which were later aliquoted and stored at −20 °C for no longer than 1 month. The following compounds were used: cariprazine hydrochloride (CAS No. 1083076-69-0; TargetMol); aripiprazole (CAS No. 129722-12-9; Sigma-Aldrich); and haloperidol (CAS No. 52-86-8; Sigma-Aldrich). Ready-to-use solutions were freshly prepared on the day of the experiment by diluting the 10 mM stock solutions in artificial cerebrospinal fluid (ACSF) or neuronal culture medium, with pure DMSO used to equalize final vehicle concentration (0.1% DMSO) across conditions.

#### Multielectrode array (MEA) recordings

The Maestro Edge (Axion Biosystems) multielectrode array (MEA) system was used to record network activity from cultured mouse hippocampal neurons (sampling rate: 12.5 kHz, band-pass filtering: 200-3,000 Hz) at 9-11 days *in vitro* (DIV). Neurons were cultured in 24-well CytoView MEA plates (Axion Biosystems; 16 electrodes/well) and recorded under humidified carbogen atmosphere (5% CO□) at 37 °C temperature.

Baseline firing and bursting activity patterns were recorded for 60 minutes, after which each of the three antipsychotic compounds was administered directly to a separate well. Recordings were resumed following a 20-minute stabilization period. Root mean square (RMS) noise levels in the recorded voltage traces were below 3 µV throughout all recordings.

Action potentials were detected using the Axion Navigator software’s (AxIS) adaptive threshold-crossing algorithm, with the detection threshold set to 5.2 x the standard deviation (SD) of the signal. The percentage of active electrodes exceeded 50% in all wells and was used as a quality control criterion.

Spike times and latencies were analyzed across channels using the NeuroExpress (NEx) software package (developed by A.Sz). Network bursts were defined as clusters of spikes separated by intraburst interspike intervals [(ISIs) ≤ 0.4 s] identified by the NEx software from spike lists simultaneously generated with spike detection by the AxIS software. Network-wide synchronization of firing activity was quantified by calculating Pearson’s correlation coefficient matrices derived from spike density functions of individual spike trains. Spike density functions were calculated by convolving the spike arrival time sequences with a unit-area, 0.5 s half-width Gaussian function (kernel), at 10 sample/s resolution.

#### Whole-cell patch clamp recordings

Whole-cell patch clamp recordings were performed on primary hippocampal cell cultures bathed in ACSF containing 140 mM NaCl, 5 mM KCl, 2 mM CaCl_2_, 1 mM MgCl_2_, 5 mM HEPES, and 10 mM D-glucose (pH adjusted to 7.4). Microelectrodes were pulled from 1B150F-4 borosilicate glass capillaries (World Precision Instruments) using a P-97 Flaming/Brown micropipette puller (Sutter Instruments) and filled with intracellular solution containing 100 mM K-gluconate, 10 mM KCl, 20 mM KOH, 2 mM MgCl_2_, 2 mM NaCl, 10 mM HEPES, 0.2 mM EGTA, and 5 mM D-glucose.

Electrode resistance ranged between 4-6 MΩ, and access resistance after membrane break-in was typically <12 MΩ and stable during recordings. Neurons were visualized using a Scientifica SliceScope microscope equipped with an Olympus UMPLFLN 20x water-immersion objective, and QImaging camera controlled by QCapture software.

Current-clamp recordings were performed using a stepwise current-injection protocol, consisting of 400-ms hyperpolarizing and depolarizing current steps, starting from −150 pA up to +200 pA with 5 pA increments, to evoke voltage responses from patched neurons. When indicated by the initial current-step responses (e.g. pronounced outward rectification or the presence of spikelets), additional targeted protocols were applied.

Signals were amplified using a MultiClamp 700B amplifier (Molecular Devices) and digitized at 20 kHz using a National Instruments PCIe-6220 data acquisition board, with data acquired using DASYLab 2016 software (Windows 11). Recorded traces were analyzed using NEx.

#### Computational modeling

To aid the interpretation of the cariprazine-induced effects on firing properties we constructed 3-compartmental model neurons based on the Hodgkin-Huxley formalism, and implemented those in our custom-made modeling platform SNMod (23, 24). The first canonical model was designed to reproduce the physiological properties of regular firing and stuttering neuronal subtypes in hippocampal cultures. It featured a fast Na-current, a delayed rectifying K-current, a hyperpolarization-activated cation current (Ih), a slowly activating M-type K-current and a strong D-type K-current. The second model, designed to reproduce the physiological properties of delayed firing type hippocampal neurons, was similar to the first one, but the h-current was replaced with an inward rectifying K-current here. The voltage-dependent membrane currents were distributed across the somatic, axonal and dendritic compartments as shown in Supplementary Table 7. Physiological parameters such as membrane resistance, voltage sag and rheobase were carefully adjusted using experimental data to attain the best match between simulated and recorded voltage traces. All intrinsic voltage-dependent currents were calculated according to the following equation

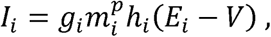

where *i* represents the individual current type, *g_i_* is the maximal conductance of the current, *m_i_* is the activation variable, *p* is the exponent of the activation term, *h_i_* is the inactivation variable (either first-order or absent), and *E_i_* is the reversal potential. Activation (*m*) and inactivation (*h*) variables were calculated from the equation

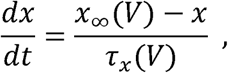

where *x* is either *m* or *h*, and voltage-dependent steady-state activation and inactivation were described by the sigmoid function

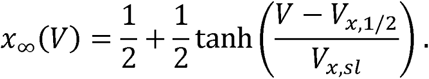

The midpoint *V_x,1/2_* and slope *V_x,sl_* parameters of the sigmoids and the other kinetic parameters are shown in Supplementary Table 7. Time constants of the activation and inactivation were bell-shaped functions of the membrane potential according to the equation:

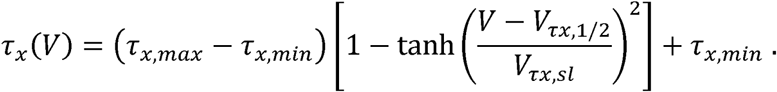

Here, *t_x,max_* and *t_x,min_* indicate the maximal and minimal values of the bell-shaped functions, *V_tx,1/2_* indicates their midpoint and *V_tx,sl_* sets the slope of the functions.

#### Reverse transcription quantitative real-time PCR (RT-qPCR)

Total RNA was isolated using RNeasy Mini or Micro Kit (Qiagen) following the manufacturer’s instructions. Complementary DNA (cDNA) was synthesized from purified RNA using Maxima First Strand cDNA Synthesis Kit (Thermo Fisher Scientific). Reverse transcription quantitative real-time (RT-qPCR) reactions were prepared by combining gene-specific primers with LightCycler 480 SYBR Green I Master mix (Roche) and were run on a Bio-Rad CFX Opus System.

Gene expression levels were normalized to two mouse reference genes, *PARP* and *RSP29*, and relative expression values were calculated using the ΔΔCt method.

**Table 1.**
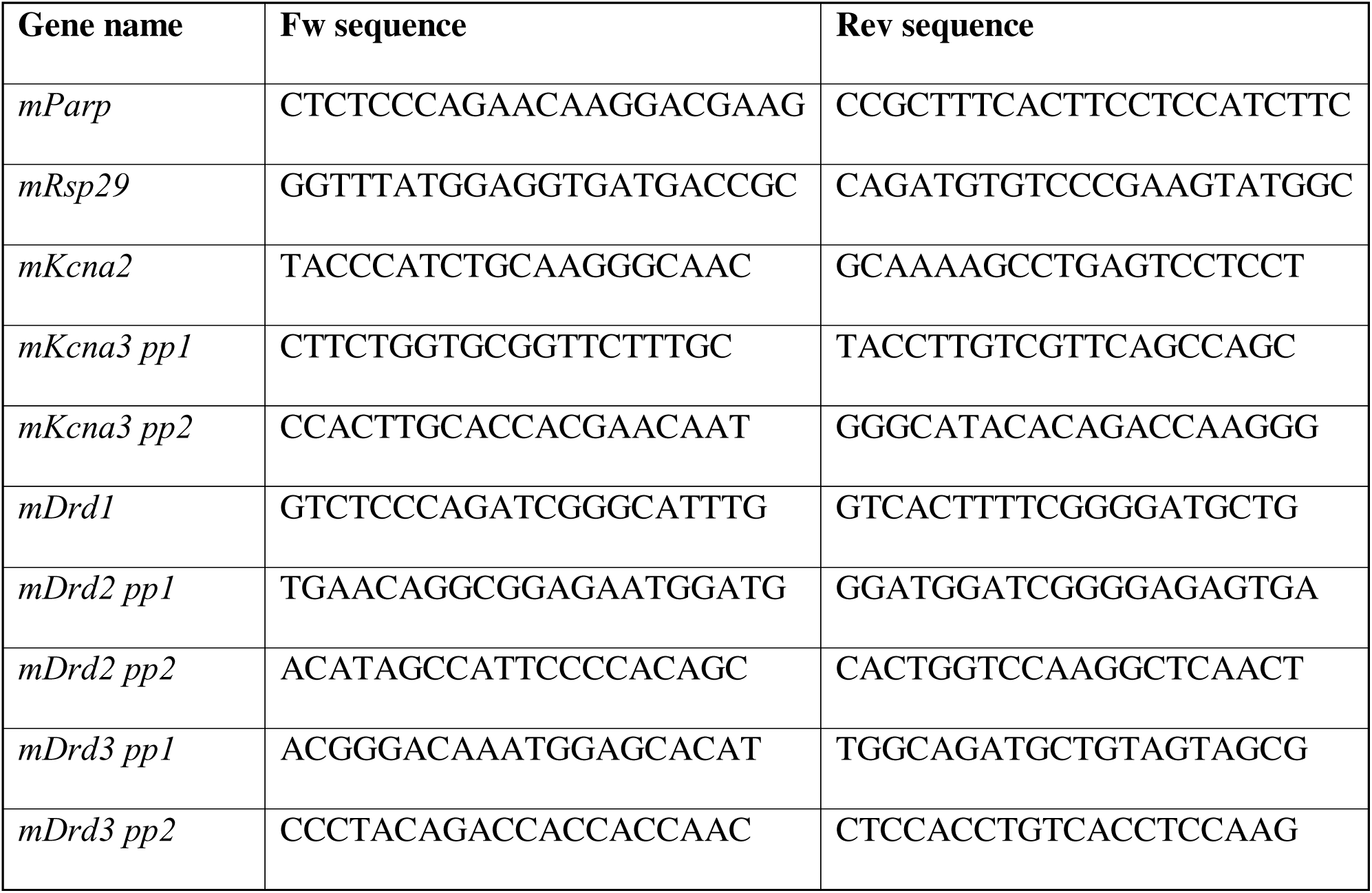
Primer sequences used for RT-qPCR.

#### Viability Assay

CD-1 hippocampal neurons were seeded into 96-well plates at a density of 13,000 cell/well. Cells were exposed to cariprazine (cat. no. T8382) across a concentration range of 0.03, 0.1, 0.3, 1, 3, and 10 µM. Two treatment paradigms were applied: an acute exposure lasting 1.5 hours and a chronic exposure lasting 72 hours.

Cariprazine stock solutions were prepared in DMSO as specified above and subsequently diluted in culture medium to obtain the final working concentrations. Vehicle control wells received an equivalent volume of DMSO to account for solvent-specific effects. Cell viability was assessed using the CellTiter-Glo Luminescent Cell Viability Assay (Promega) according to the manufacturer’s instructions. Luminescent signals were recorded with a Clariostar Flash multimode plate reader (BMG Labtech).

Viability values were normalized to untreated control wells, defined as 100% viability, while water-treated wells were used as the 0% viability reference. Data analysis was performed using Prism 10 software (GraphPad).

#### Immunocytochemistry (ICC)

Cells were fixed with 4% paraformaldehyde (PFA) for 10 minutes, followed by permeabilization with 0.1% Triton X-100 in PBS (Gibco, Thermo Fisher Scientific) for 10 minutes. Cells were then blocked using donkey serum diluted in PBS (50 μL serum per 1 mL PBS) for 30 minutes. Cultures were incubated with anti-chicken Map2 (RRID: AB_2138153, Cat# ab5392, Abcam) and anti-rabbit Kv1.2 (RRID: AB_2545562, Cat# PA5-144086, Thermo Fisher Scientific) primary antibodies overnight at 4 °C, diluted 1:20,000 (Map2) and 2 μg/mL (Kv1.2) in blocking solution. On the following day, cells were washed twice with PBS and incubated with secondary antibodies Cy™3 AffiniPure Donkey Anti-Chicken IgY (IgG) (H+L) and Cy™5 AffiniPure Donkey Anti-Rabbit IgG (H+L) (Jackson ImmunoResearch), diluted 1:200 in blocking solution, for 2 hours at room temperature in the dark. Nuclei were counterstained with DAPI for 15 minutes (1:1,000), followed by PBS washes. Wells were filled with 300 μL of PBS, sealed, and stored at 4°C in the dark until imaging. High-content automated imaging and analysis was performed using the CellInsight CX5 High-Content Screening platform (Thermo Scientific). Additional cultures were stained for Tau (RRID: AB_2886280, Cat# GTX130462, GeneTex) and Map2 alongside with DAPI to define neuronal populations in mixed neuron-glia cultures.

#### High-content automated microscopy (HCA)

High-content automated microscopy was performed using the CellInsight CX5 High-Content Screening platform (Thermo Scientific), following as previously described (25). Immunostained cultures were imaged at 10x magnification using automated acquisition, and quantitative analysis was conducted using the Cell Compartment Analysis (CCA) module of the CX5 HCS software.

Nuclei (DAPI□) and neuronal cells (Map2□) were identified based on fluorescence intensity, object area, and shape parameters. Objects with abnormal morphology, extreme intensity values, or partial segmentation at image borders were excluded to ensure robust cell identification. Kv1.2 signal was quantified within Map2 neuronal compartments using intensity-based thresholds defined from representative control images.

For each condition, multiple (169) fields per well were analyzed, and only wells containing a minimum number of valid neurons were included in downstream analyses. Higher-magnification (20x) images were acquired for qualitative visualization of Map2 and Kv1.2 subcellular localization. All imaging and analysis parameters were kept constant across experimental conditions.

#### Statistical analysis

Electrophysiological datasets. Single-unit recordings from the same neurons were compared using Wilcoxon paired-sample signed-rank test to evaluate changes between control and treatment conditions. Multiunit activity was compared across the most active 4 electrodes selected for each well in baseline activity measurements, and the same electrodes were analyzed between control and treatment conditions. Within cultures, activity was correlated for selected electrode pairs with the Pearson’s correlation method and parameter differences were evaluated by using Friedman ANOVA with post-hoc tests (Dunn’s test, Wilcoxon-Nemenyi-McDonald-Thompson test).

Immunocytochemistry and real-time quantitative PCR datasets. Kruskal-Wallis tests followed by Dunn’s post-hoc comparisons were used to assess statistical differences in Kv1.2 abundance and *Kcna* expression between control and treated cell cultures. For *Drd* expression data, we used Welch’s unpaired two-sample t-tests. Statistical analysis was performed using Prism10 (GraphPad) and OriginPro (OriginLab).

## RESULTS

### Real-world clinical outcomes associated with D2/D3 partial agonist antipsychotics in schizophrenia

We analyzed nationwide real-world healthcare data from patients with schizophrenia-spectrum disorders receiving antipsychotic medications acting primarily at dopamine D2/D3 receptors. The analysis focused on haloperidol (first-generation antipsychotic; D2 antagonist), aripiprazole (third-generation antipsychotic; D2/D3 partial agonist), and cariprazine (third-generation antipsychotic; D3-preferring D2/D3 partial agonist), reflecting both their shared dopaminergic targets and their distinct pharmacological and clinical profiles (Fig. 1A). The unmatched study cohort consisted of 12,166 patients who met all cohort inclusion criteria. Specifically, patients were required i) to have received only one of the three study antipsychotics during the index period (between 1^st^ January 2018 and 31^st^ December 2018), ii) to be alive at the end of the index period, and iii) to have no pregnancy-related or iv) substance-use-specific diagnostic codes recorded by the end of the index period. The cohort included 1,573 patients receiving cariprazine, 6,415 receiving aripiprazole, and 4,178 receiving haloperidol (Supplementary Table 1). The baseline cohort characteristics differed substantially across treatment groups. Patients receiving haloperidol were generally older and had a higher burden of several somatic and neurological comorbidities, including dementia, epilepsy, heart failure, cerebrovascular disease, and peripheral vascular disease. In contrast, patients receiving cariprazine or aripiprazole more frequently had recorded affective or anxiety-related psychiatric comorbidities and more frequent concomitant antidepressant use.

**Figure 1.**
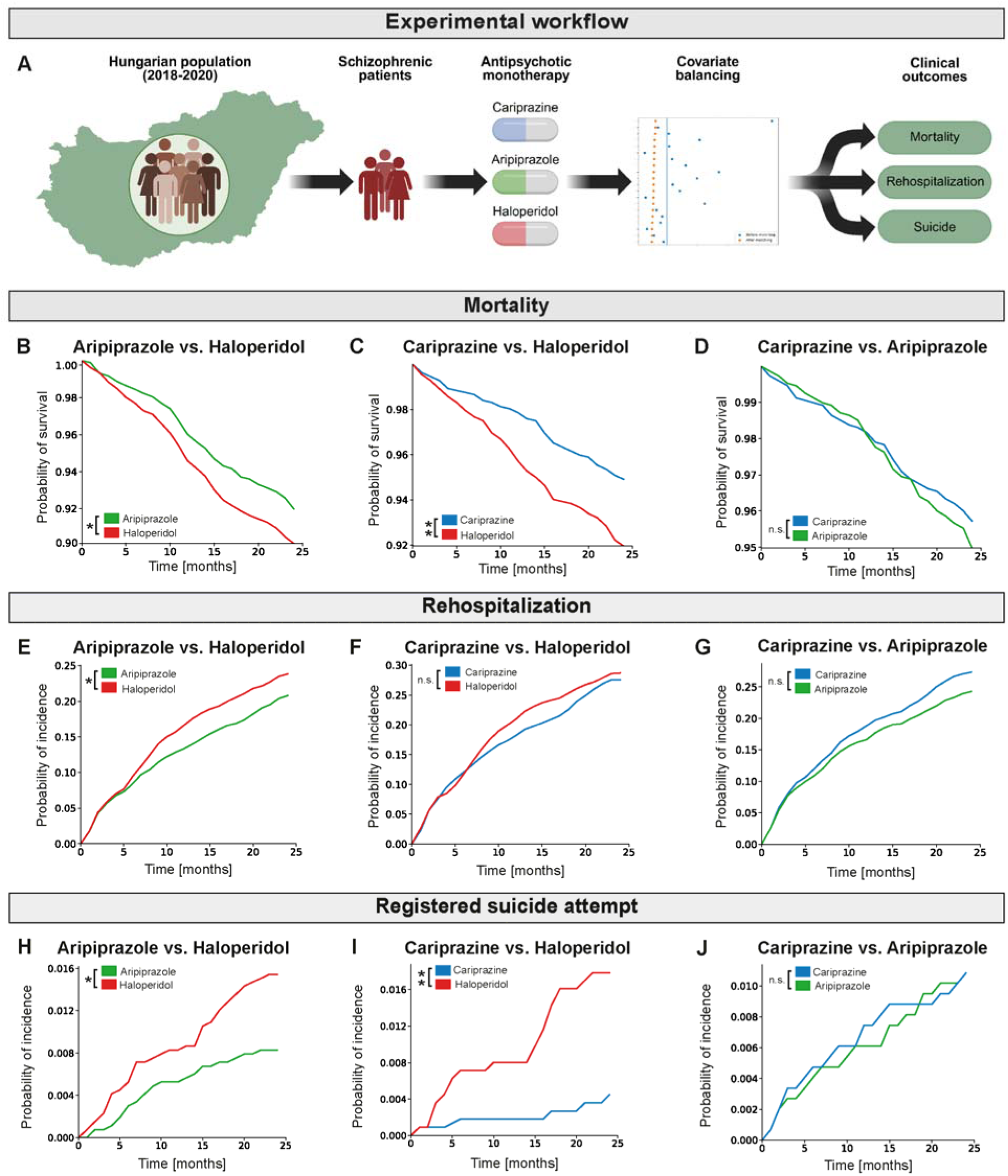
Clinical outcome probabilities after index-drug monotherapy in patients with schizophrenia-spectrum disorder. (A) Overview of the real-world study workflow. Hungarian patients with schizophrenia-spectrum disorders receiving cariprazine, aripiprazole, or haloperidol were identified from nationwide healthcare data, matched within sex and 5-year age strata using high-dimensional propensity score matching, and analyzed for mortality, rehospitalization, and first registered suicide attempts. Matching improved balance among propensity score model-included features and improved propensity score overlap between comparison groups. (B-D) Overall survival probability. (E-G) Cumulative incidence of first psychiatric rehospitalization. (H-J) Cumulative incidence of first registered suicide attempt. For each outcome, pairwise comparisons are shown for aripiprazole versus haloperidol (B, E, H), cariprazine versus haloperidol (C, F, I), and cariprazine versus aripiprazole (D, G, J). Matched cohort sizes were as follows: aripiprazole vs haloperidol, n=2,656 pairs; cariprazine vs haloperidol, n=1,119 pairs; cariprazine vs aripiprazole, n=1,475 pairs. *p<0.05; pairwise log-rank tests were used for overall survival, and pairwise Gray’s tests were used for first psychiatric rehospitalization and first registered suicide attempt. Exact p-values are provided in the main text.

To reduce baseline confounding, pairwise high-dimensional propensity score matching was performed separately for each drug comparison. The three matched cohorts included 1,475 cariprazine-treated and 1,475 aripiprazole-treated patients, 1,119 cariprazine-treated and 1,119 haloperidol-treated patients, and 2,656 aripiprazole-treated and 2,656 haloperidol-treated patients. Overall balance among variables included in the propensity score model improved substantially after matching, and propensity score overlap between treatment groups was improved (Supplementary Fig. 1). Median post-matching standardized mean differences for model-included variables were low across comparisons: 0.018 for cariprazine–aripiprazole, 0.032 for cariprazine–haloperidol, and 0.027 for aripiprazole–haloperidol (Supplementary Table 2). Some clinically relevant variables reported in Supplementary Table 1 were not fully represented in the final matching feature set, these variables were evaluated separately when assessing residual imbalance (Supplementary Table 2).

Although post-matching balance was adequate for variables included in the propensity score model, separate assessment of selected clinically relevant baseline characteristics revealed limited residual imbalance in some comparisons. In the cariprazine–aripiprazole matched cohort, F20 schizophrenia diagnosis remained more frequent among cariprazine-treated patients, with a post-matching SMD of 0.168, whereas lithium use was more frequent among aripiprazole-treated patients, with a post-matching SMD of −0.110. In the cariprazine–haloperidol matched cohort, antidepressant use and F20 schizophrenia diagnosis remained more frequent among cariprazine-treated patients, with post-matching SMDs of 0.117 and 0.118, respectively, while lithium use was more frequent among haloperidol-treated patients, with a post-matching SMD of −0.173. In the aripiprazole–haloperidol matched cohort, no assessed clinically relevant baseline characteristic exceeded the conventional absolute SMD threshold of 0.1. These findings indicate that residual measured confounding cannot be fully excluded, particularly in comparisons involving cariprazine.

Across the matched cohorts, patients receiving haloperidol generally showed less favorable clinical outcomes than those receiving aripiprazole or cariprazine, particularly for all-cause mortality and first registered suicide attempt. Kaplan-Meier survival analyses showed significantly lower overall survival probabilities in the haloperidol group than in the aripiprazole and cariprazine groups (Fig. 1B-D). The difference was statistically significant for both aripiprazole versus haloperidol (p = 1.65 × 10^-2; Fig. 1B) and cariprazine versus haloperidol (p = 4.70 × 10^-3; Fig. 1C). In contrast, survival probabilities for cariprazine and aripiprazole largely overlapped, with no statistically significant difference between the two partial agonists (p = 3.43 × 10^-1; Fig. 1D).

Psychiatric rehospitalization, used as a proxy for relapse or clinical destabilization, showed a less consistent pattern across pairwise comparisons than all-cause mortality (Fig. 1E–G). In the aripiprazole-haloperidol comparison, haloperidol-treated patients had a significantly higher cumulative incidence of first psychiatric rehospitalization (p = 1.28 × 10^-2; Fig. 1E). The comparison between cariprazine and haloperidol did not reach statistical significance (p = 6.06 × 10^-1; Fig. 1F), and no statistically significant difference was observed between cariprazine and aripiprazole (p = 5.39 × 10^-2; Fig 1G).

For first registered suicide attempt, both comparisons with haloperidol showed lower cumulative incidence among patients receiving aripiprazole or cariprazine (Fig. 1H-J). The cumulative incidence differed significantly between aripiprazole and haloperidol (p = 2.36 × 10^-2; Fig. 1H) and between cariprazine and haloperidol (p = 2.54 × 10^-3; Fig. 1I), whereas no significant difference was observed between cariprazine and aripiprazole (p = 8.61 × 10^-1; Fig. 1J). However, because the number of first registered suicide attempt events was low in the matched cohorts, these results should be interpreted with caution.

Taken together, these real-world findings suggest that aripiprazole and cariprazine, the two antipsychotics with D2/D3 partial agonist activity examined in this study, were associated with more favorable survival outcomes than haloperidol in patients with schizophrenia-spectrum disorders. Similar patterns were observed for first registered suicide attempts, although event counts were low. For psychiatric rehospitalization, aripiprazole was associated with a significantly lower cumulative incidence than haloperidol, whereas cariprazine did not differ significantly from either haloperidol or aripiprazole. Across all examined outcomes, no statistically significant differences were observed between cariprazine and aripiprazole.

### Consistent network bursting is observed in cultured hippocampal neuron networks

To bridge these clinical observations with a more mechanistic understanding of antipsychotic action, we next employed well-characterized primary hippocampal cultures from CD-1 mice to investigate the effects of the previously described antipsychotics at the network and cellular levels using multielectrode array (MEA) and patch clamp electrophysiology. To validate the cellular composition of the cultures, we first performed immunocytochemical (ICC) labeling combined with high-content automated microscopy (HCA) analysis. Quantification of Tau and Map2 cells revealed that approximately 15% of DAPI cells consistently exhibited neuronal marker expression across cultures, with a substantial overlap between Tau and Map2 labeling, in line with previous reports on embryonic mouse hippocampal cultures (Supplementary Fig. 2B). This confirmed the presence of a stable neuronal population suitable for functional network and single-cell electrophysiological recordings.

To assess the relevance of dopaminergic signaling in this *in vitro* system, we next examined dopamine receptor expression using RT-qPCR. We detected clear and robust expression of *Drd1*, *Drd2*, and *Drd3* transcripts in hippocampal cultures, supporting the suitability of this model for investigating dopamine receptor-dependent effects of antipsychotic compounds (Supplementary Fig 2C).

In addition, to define concentration ranges suitable for electrophysiological experiments, we assessed cariprazine cytotoxicity using a luminescence-based cell viability assay under both acute (1.5 h) and chronic (72 h) exposure conditions. Based on these measurements, non-cytotoxic concentration ranges were selected for subsequent MEA and patch clamp experiments. Concentrations of haloperidol and aripiprazole were chosen based on previously published in vitro studies and were matched to cariprazine treatments to enable direct pharmacological comparison across compounds (26, 27).

To obtain an initial functional assessment of how cariprazine and other antipsychotic drugs influence neuronal network dynamics, we employed the Maestro Edge MEA system to record spontaneous activity from primary mouse hippocampal neuron cultures between DIV9 and DIV11. This platform enables long-term, non-invasive recordings from 24 independent cultures (wells), each monitored by 16 extracellular electrodes, allowing parallel assessment of network-level firing patterns.

For each well, spontaneous activity was first recorded under baseline conditions for 60 minutes, followed by recordings up to 2 hours after acute application of antipsychotic compounds at final concentration of 1 µM and 10 µM.

Under baseline conditions, well-developed hippocampal cultures displayed robust spontaneous firing activity characterized by synchronous network bursts and transient oscillatory patterns, such as theta rhythm activity (Fig. 2B and C). Although the overall temporal structure of bursting was consistent across cultures, network activity remained highly variable and complex, often interrupted by silent periods lasting up to 2 minutes. This intrinsic variability warranted to monitor the network activity for extended durations to reliably assess both frequent short-lasting bursts and more sporadically occurring large-scale network events.

**Figure 2.**
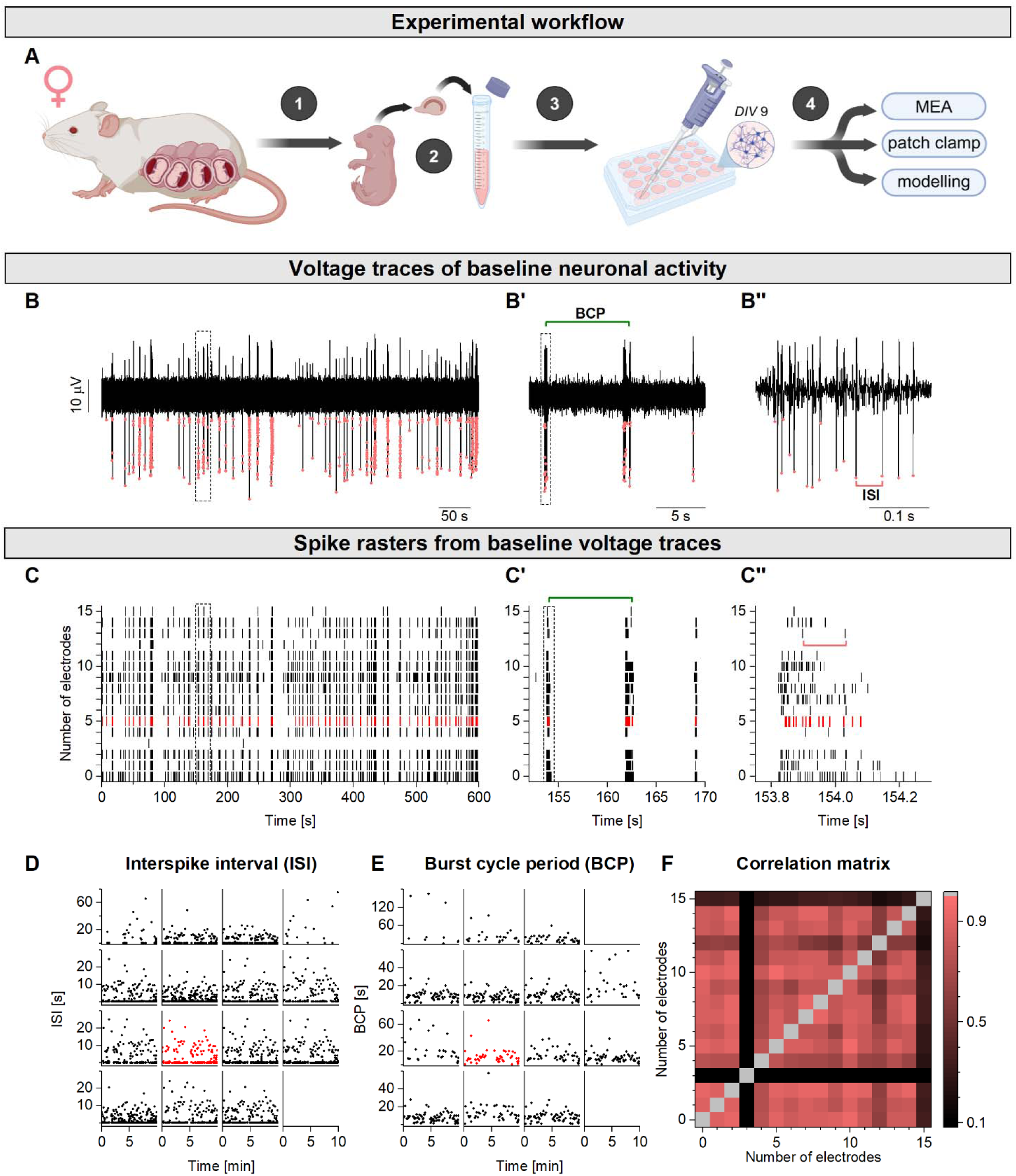
Multielectrode array recordings of cultured hippocampal neurons reveal robust bursting activity and network-wide synchronization. (A) Schematic overview of the experimental workflow using primary CD1 mouse hippocampal neurons. (B) Representative voltage trace recorded from a single MEA electrode containing spikes of variable amplitude, likely originating from two distinct neuronal units. Red dots indicate detected spikes. Scale bar: 10 μV; 50 s. (B’) Enlarged segment illustrating burst cycle periods (BCP). Scale bar: 5 s. (B’’) Magnified segment highlighting intraburst interspike intervals (ISI). Scale bar: 0.1 s. (C) Spike raster plot from 16 electrodes recorded over 600 s. Red ticks correspond to spikes shown in B. (C’) Enlarged raster segment illustrating burst timing and BCP. (C’’) High-resolution view of intraburst ISIs. (D) Interspike intervals plotted as a function of elapsed time for the 16 electrodes arranged in a 4 x 4 grid. (E) Burst cycle periods. (F) Pairwise Pearson’s correlation coefficients calculated from spike density functions of individual electrodes, reflecting strong network-level synchronization.

Baseline firing patterns from three representative cultures are shown as raster plots with corresponding spike density functions (Fig. 3). In these plots, ticks of variable thickness represent bursts typically composed of multiple spikes. While all cultures exhibited pronounced and reproducible network bursting, substantial variability was evident at shorter timescales (minutes). In addition to regular network bursts, highly synchronized, prolonged burst episodes (“superbursts”) were observed in several cultures (Fig. 3A, G and M), recurring with longer cycle periods of approximately 2-5 minutes.

**Figure 3.**
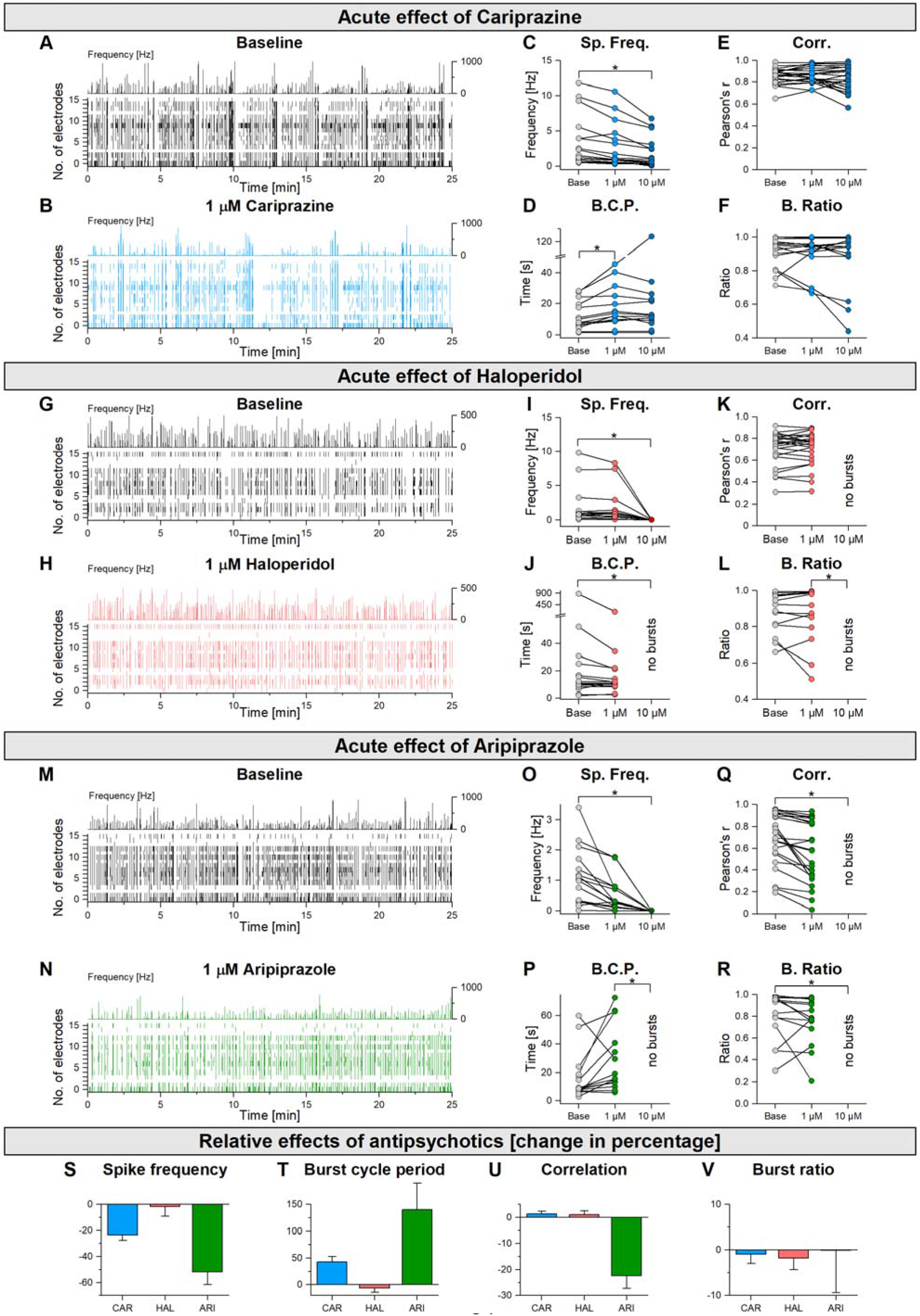
Antipsychotics reduce the overall firing activity of cultured hippocampal neurons and induce variable degree of reduction of network synchronization. (A) Spike raster plot of a 30 min baseline MEA recording (16 electrodes) with the corresponding total spike density function (upper graph). (B) Raster plot of the same well following 1 µM cariprazine application reveals a reduction of firing activity and prolonged silent periods between bursts. (C) Dose-dependence of the CAR effect on total firing rate. (D) Burst cycle periods and (E) pairwise Pearson’s correlation coefficients. (F) Burst ratio was calculated for each electrode as the ratio of the number of spikes within bursts to the total number of spikes. (G) Raster plot with spike density function in control and (H) following haloperidol application. (I-L) Corresponding dose dependence plots for the 3 parameters. BCPs and correlations are not available when 10 µM HAL entirely suppressed the firing activity. (M and N) Raster plots in control and after aripiprazole application. (O-R) Corresponding dose dependence plots for aripiprazole. Data are shown as mean ± SEM. Each dot represents an electrode. For quantitative comparisons, the same four electrodes with the highest baseline activity were analyzed before and after treatment in each well. Statistical significance was assessed using Friedman ANOVA [ns, p ≥ 0.05; *p < 0.05; **p < 0.01]. n = 4 electrodes/well, 4 wells/condition.

### Acute antipsychotic application modulates hippocampal network activity and synchronization in a drug-dependent manner

To minimize non-specific changes of network activity, antipsychotic compounds were applied by direct addition of small-volume, concentrated solutions into the wells, rather than by large-scale medium replacement. This approach was motivated by prior observations showing that routine 50% medium exchange caused a pronounced and prolonged perturbation in network activity. In such control recordings, firing activity was markedly reduced and burst cycle periods were prolonged for 40-60 minutes, followed by gradual recovery to baseline levels (Supplementary Fig. 3). These transient effects likely reflect changes in osmolarity, pH, or other physio-chemical parameters of the extracellular environment. Based on these observations, drugs were administered by adding 0.6 µl of concentrated stock solution directly to the wells, yielding final concentrations of 1 μM and 10 μM. Firing activity of hippocampal neuron cultures was recorded before and after drug application, generating time series of 60-minute duration. Figure 3 illustrates the firing patterns from four representative cultures under control conditions and following application of antipsychotic drugs (1 µM). For each well, 4 out of 16 electrodes exhibiting the most robust baseline activity were selected for analysis per condition. From these spike trains, mean firing rate, mean burst cycle period, and pairwise correlation coefficients were calculated.

Cariprazine decreased firing activity across all four cultures. Specifically, 1 µM and 10 µM cariprazine decreased the overall spike frequency by 23.3 ± 4.1 % and 50.1 ± 3.2 %, respectively (Fig. 3C, S), while burst cycle periods increased by 42.6 ± 10.1 % and 57.0 ± 19.8 % (Fig. 3D, T). Despite these pronounced changes in firing and burst timing, network synchronization was not affected by the treatment as shown by the consistently high values of the correlation coefficients calculated from the spike density functions of individual electrodes (Fig. 3E). This suggests that acute cariprazine application primarily modulates intrinsic neuronal excitability rather than disrupting synaptic coupling within the network.

In contrast, aripiprazole produced a stronger suppression of firing activity, reducing firing rates by 52.5 ± 6.4% and 51.7 ± 9.8% at 1 µM and 10 µM, respectively (Fig. 3O, S). At 10 µM, cariprazine further reduced the firing rate, whereas haloperidol and aripiprazole effectively blocked all spiking activity in the cultures (Fig. 3I and O). Notably, aripiprazole - but not haloperidol - also reduced network synchronization, as shown by pairwise correlation coefficients (Fig. 3K and Q). Together, these observations suggest that cariprazine and haloperidol exert more similar effects on hippocampal network organization, potentially acting predominantly through modulation of intrinsic cellular properties rather than synaptic connectivity in this *in vitro* system.

Washout of the antipsychotic compounds restored the network activity towards baseline levels (Supplementary Fig. 4). Although the high concentrations applied during the second recording phase completely suppressed bursting, robust and synchronous network activity was consistently recovered following washout in all cultures (Supplementary Fig. 4B-C, E-F, H-I). As observed previously, complete medium replacement during washout transiently altered firing patterns, consistent with the effects seen during medium exchange controls (Supplementary Fig. 3B and E).

### Intrinsic excitability of hippocampal neurons is decreased by cariprazine

The MEA experiments indicated that cariprazine reduces overall firing activity and prolongs burst intervals in hippocampal networks without disrupting network synchronization. This pattern suggests that cariprazine may act by altering the intrinsic excitability of the neurons, which is primarily regulated by voltage-dependent membrane currents. Homeostatic up- or downregulation of such currents, developing on longer timescales (24 - 48 hours) and targeting specific ion channels, has been reported previously in primary neuronal cultures using combined MEA and patch clamp techniques (28, 29). We therefore next aimed to identify potential membrane-level targets of cariprazine using whole-cell patch clamp recordings and current step stimulation. This allows detailed characterization of both passive and active membrane properties, many of which are closely associated to specific voltage-gated ion channels.

Under control conditions, nearly two-thirds of neurons in primary hippocampal cultures showed a regular action potential firing phenotype (Fig. 4M), characterized by spike frequency adaptation and a moderate voltage sag (Fig. 4A) under hyperpolarizing current steps, consistent with the presence of the h-current. Another, smaller subset of neurons featured lower intrinsic excitability, a characteristic “stuttering” firing pattern or just a single action potential emitted during depolarizing current steps. The action potentials in such neurons exhibited a pronounced spike afterhyperpolarization, a phenomenon commonly associated with fast-spiking interneurons of the hippocampus. Finally, a distinct population of neurons exhibited robust K_ir_-current-mediated inward rectification together with regular firing output - these cells, referred to as delayed firing types, could be readily distinguished from the other phenotypes by their reduced membrane resistance and lack of voltage sag.

**Figure 4.**
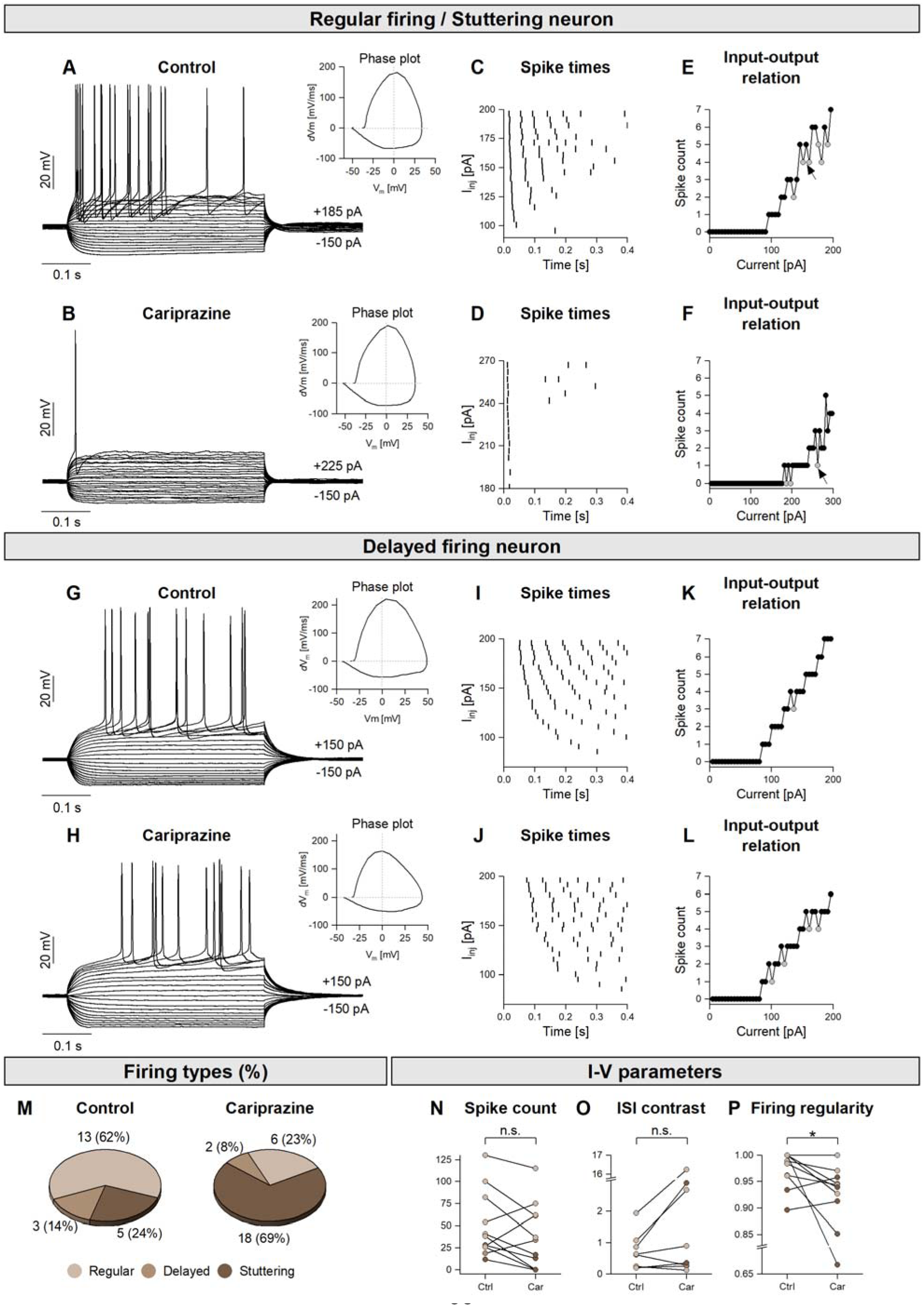
Acute application of cariprazine reduces the intrinsic excitability of hippocampal neurons. (A) Voltage responses of a regular-firing neuron stimulated by stepwise current injections (from −150 to +185 pA) under control conditions. (B) Voltage responses of the same neuron following application of 1 µM cariprazine, showing single-spike firing up to +225 pA. (C and D) Peri-stimulus spike raster plots recorded under control conditions (C) and after cariprazine treatment (D). (E and F) Spike count as a function of injected current under control conditions (E) and after cariprazine application (F). Light grey symbols indicate traces at which spike counts temporally deviate from a monotonically increasing trend, reflecting a stuttering firing behavior. (G and H) Voltage responses of a delayed-firing neuron under control conditions (G) and following cariprazine application (H), demonstrating reduced sensitivity to the drug. (I and J) Peri-stimulus spike raster plots of the delayed-firing neuron under control conditions (I) and after cariprazine application (J). (K and L) Spike count–current relationships for the delayed-firing neuron in control conditions (K) and in the presence of cariprazine (L). Firing becomes less regular and reduced after cariprazine application relative to the control. (M) Distribution of firing phenotypes in 21 hippocampal neurons recorded under control conditions, with regular-firing neurons being the most abundant. Phenotype distribution following cariprazine exposure, showing an increased proportion of stuttering neurons. (N) Cumulative spike counts in control conditions and following cariprazine application. (O) Interspike interval (ISI) contrast before and after cariprazine application. (P) Quantification of firing regularity. (N, O, P) Each dot corresponds to a single neuron. Single-unit parameter changes were evaluated using Wilcoxon paired-sample signed-rank tests.

Acute exposure of hippocampal neurons to 1 µM cariprazine reduced their firing output during depolarizing current steps (Fig. 4A-F; spike count decrease: 24.6 ± 23.1%, n=11) without significantly changing their membrane resistance or voltage sag. Regular firing or stuttering neurons displayed the strongest reduction in firing (Fig. 4A and B), whereas delayed-firing neurons were less sensitive to cariprazine (Fig. 4G-L).

In addition, cariprazine significantly altered firing regularity (Fig. 4P) as shown by an increased variability of ISIs under suprathreshold current steps. To quantify this effect, we introduced a metric called ISI contrast. ISI contrast increased following cariprazine application (272 ± 100%, n=7; Fig. 4O), and this increase coincided with the reduction in intrinsic excitability observed in the same neurons (Fig. 4N).

Beyond neurons recorded before and during acute wash-in of the drug, we also studied cells that had been already exposed to cariprazine for up to 1-hour prior to recording. In these neurons, stuttering firing responses and low intrinsic excitability were detected, substantially more frequently than under control conditions (Fig. 4M). Quantitatively, the proportion of stuttering type neurons nearly tripled, indicating a population-level shift in the intrinsic firing properties induced by cariprazine treatment.

Together, these single-cell experiments revealed cell-type-dependent reductions in the intrinsic excitability and firing regularity of neurons in primary hippocampal cultures. These changes are consistent with upregulation of specific voltage-dependent potassium currents. In particular, Kv1 channel-mediated, slowly inactivating D-type potassium currents are known to facilitate stuttering firing behavior and reduce overall firing output (24, 30). Based on these observations, we next investigated whether modulation of the D-current could account for the effects of cariprazine using a single-cell computational model.

### Cariprazine-induced effects investigated in a computational model

Based on our current patch clamp data and extensive prior observations on the physiological properties of cultured hippocampal neurons, we created two types of canonical conductance-based neuron models representing the firing phenotypes observed experimentally. The regular-firing model featured an h-current, a non-inactivating M-type potassium-current, and a slowly inactivating D-type potassium-current, in addition to spike-generating Na- and delayed rectifier potassium (K_dr_) currents. This model exhibited voltage sag under hyperpolarizing current steps (Fig. 5A) and a monotonically increasing number of spikes in response to increasing depolarizing current steps (Fig. 5C). The model closely reproduced many of the passive and active membrane properties of the biological neurons, as measured using patch clamp.

**Figure 5.**
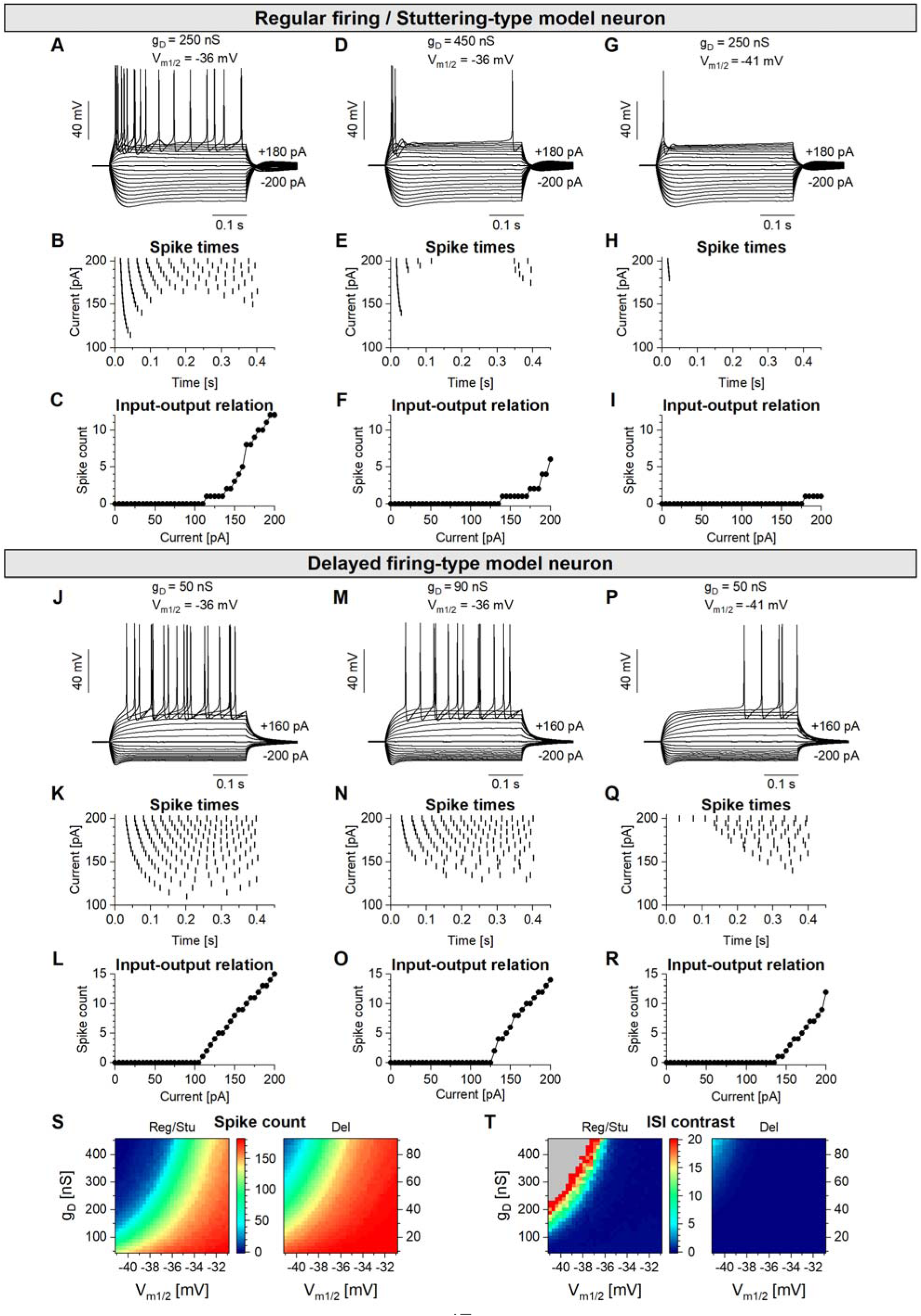
Manipulation of the D-type potassium current differentially regulates intrinsic excitability and firing patterns in model neurons. (A) Voltage responses of the regular-firing type model neuron under stepwise somatic current injections. (B) Peri-stimulus spike raster plot and (C) input-output (I-O) relationship of the canonical regular-firing model. (D) Voltage responses of the same model neuron after incrementing by +80% of the maximal conductance of its D-current. (E) Corresponding spike raster plot and (F) I-O relationship, where after the maximal D-current conductance is 450 nS. (G) Voltage responses of the regular-firing model neuron with the original maximal conductance (250 nS) but after shifting the half-activation voltage of the D-current by −5 mV. (H) Raster plot and (I) I-O relationship after shifting the half-activation voltage. (J) Voltage responses of the canonical delayed-firing model neuron. (K) Peri-stimulus spike raster plots and (L) corresponding I–O functions for the delayed-firing model in control conditions. (M) Voltage responses of the delayed-firing model after increasing the maximal conductance of the D-current by +80% to 90 nS or (P) performing a −5 mV shift of the half-activation voltage without changing the maximal conductance of D-current. (N, O, Q, R) Spike rasters and I-O functions corresponding to conductance and half-activation manipulations in the delayed firing type neuronal model. (S) Heatmap of firing intensity for the regular and delayed firing neuronal models. The cumulative spike count is plotted against the maximal conductance of the D-current and its half-activation threshold (33 x 33 parameter combinations). (T) Heatmap of ISI contrast values demonstrating firing regularity for the regular- and delayed firing models. The grey area represents firing responses with too few spikes for the calculation of contrast values. Average interspike interval coefficient of variation plotted against the maximal conductance and half-activation voltage of the D-current (33 x 33 parameter combinations).

The delayed-firing model featured a K_ir_-current, an M-current and a D-current with a maximal conductance (g_D_=50 nS) corresponding to 20% of that used in regular firing model. This model displayed prominent inward rectification during negative current pulses (Fig. 5J) and a prolonged latency to the first spike (Fig. 5K), resulting from deactivation of the intrinsic K_ir_-current under depolarizing inputs. Despite this delayed onset, firing output remained regular, as shown by the smoothly increasing number of spikes emitted under depolarizing current steps (Fig. 5L).

Notably, cultured hippocampal neurons associated with the delayed-firing phenotype almost never displayed stuttering or outward rectification, in contrast to regular-firing neurons. This observation strongly suggests that neurons with strong intrinsic K_ir_ currents have a lower base level of D-type potassium currents than the regular-firing neurons. Consequently, we implemented a higher D-current conductance in the regular-firing model (g_D_ = 250 nS) compared to the delayed-firing model (g_D_ = 50 nS).

Increasing the maximal conductance of the D-current reduced intrinsic excitability in both models but with markedly different effects. In the regular-firing model, incrementing the D-current conductance changed the firing pattern into a more irregular one (Fig. 5B and E). Sudden decreases in the input-output (I-O) relationship (Fig. 5F) indicated stuttering behavior similar to that observed experimentally following cariprazine treatment. Importantly, this manipulation did not alter membrane resistance, time constant, or voltage sag. In contrast, the delayed-firing model was much less affected by D-current upregulation, as firing output was only slightly reduced.

In the next set of simulations, we changed the activation properties of the D-current rather than its maximal conductance. This approach was motivated by the possibility that cariprazine may alter D-current gating properties rather than increase its maximal conductance in the acute experiments. A modest leftward shift in the steady-state activation curve of the D-current – from −36 mV to −41 mV – was applied in both models. This manipulation effectively increased its potency in regulating the firing output. Half-activation points of the D-current was moved and compared to canonical firing responses. This manipulation also reduced the firing output of the neurons (Fig. 5I) and reproduced the stuttering in the first model (Fig. 5H). Again, the delayed firing type model was less affected by this manipulation than the first model (Fig. 5Q and R).

Finally, we quantified the cumulative spike counts and ISI variability across a larger set of model neurons by systematically varying both the maximal D-current conductance and its half-activation voltage in a two-dimensional grid. Current step stimulation was run to elicit the firing responses. Cumulative spike counts and ISI contrast values were then calculated for the 33 x 33 parameter arrays. By systematically varying the g_D_ and V_m1/2_ parameters of the D-current its potency was verified in regulating the intrinsic excitability (Fig. 5S) and spike timing variability (Fig. 5T) of the model neurons. Notably, ISI variability was far more sensitive to D-current manipulation in the regular-firing type of model than in the delayed-firing model.

These simulations showed that modulation of a fast-activating, slowly inactivating potassium current is sufficient to reproduce key features observed in hippocampal neurons following acute cariprazine application, including reduced excitability and increased degree of irregularity in firing. The different sensitivity of the two model phenotypes is consistent with differences in their baseline D-current contribution and is well supported with our patch clamp measurements.

### Cariprazine alters Kv1 channel distribution at the protein level without changing KCNA gene expression

Following the identification of Kv1 channel-mediated D-type potassium currents as a likely contributor to the cariprazine-induced changes in intrinsic excitability, we next examined whether prolonged cariprazine exposure alters Kv1 channel expression at the transcriptional and/or protein level.

We first examined Kv1 channel protein distribution using ICC combined with HCA. Quantification of Map2 neurons revealed that approximately 80% of cells exhibited detectable Kv1.2 immunoreactivity, and this proportion was not altered by chronic cariprazine treatment (Supplementary Fig. 5A and B). We then performed a spot-based analysis restricted to Kv1.2-positive neurons. This analysis revealed a marked redistribution of Kv1.2-positive puncta following 24 hours of cariprazine treatment compared with control conditions (Supplementary Fig. 5A-C). Notably, this effect was attenuated after 48 hours, suggesting a transient enhancement of Kv1.2 channel clustering or membrane-associated accumulation rather than a sustained increase in channel abundance.

To determine whether these protein-level changes were accompanied by transcriptional regulation, we next performed RT-qPCR analysis. No significant changes were detected in the expression of *Kcna2* or *Kcna3* following chronic cariprazine treatment compared with control cultures (Supplementary Fig. 5D and E). Likewise, transcript levels of dopamine receptor genes (*Drd1*, *Drd2*, and *Drd3*) were not significantly altered by chronic cariprazine exposure (Supplementary Fig. 5F-I).

Together, these results indicate that chronic cariprazine treatment does not induce transcriptional upregulation of Kv1 channel genes or dopamine receptors but instead selectively modulates Kv1 channel protein distribution and subcellular organization. These changes provide a molecular correlate for the observed alterations in intrinsic excitability and neuronal firing properties.

## DISCUSSION

Altered neuronal oscillations and disrupted network dynamics are well-established features of schizophrenia and related psychotic disorders, measurable by EEG and local field potential (LFP) recordings (31). Since hippocampal rhythmic activity supports learning and memory, drug-induced changes in hippocampal circuit function are likely to be particularly relevant for cognitive and negative symptom domains. Although antipsychotics can partially normalize pathological rhythmic activity, typical and atypical antipsychotics can modulate oscillations and network organization in region- and state-dependent manner (32, 33). A mechanistic understanding of how newer antipsychotics influence intrinsic excitability and network-level coordination of neuronal activity therefore remains unknown.

Here, we integrated real-world clinical outcomes with *in vitro* electrophysiology, computational modeling, and molecular analyses to define cellular mechanisms of cariprazine in hippocampal neurons. In the propensity score-matched, covariate-balanced cohorts, representing the overlapping comparison populations, exposure to haloperidol was associated with worse survival than cariprazine or aripiprazole, while psychiatric rehospitalization showed a less consistent pattern, reaching significance only in the aripiprazole–haloperidol comparison. Across MEA recordings, acute cariprazine reduced firing rate and prolonged burst intervals in a dose-dependent manner while largely preserving network synchronization, contrasting with the stronger suppression of network activity observed with haloperidol and aripiprazole under matched conditions. Whole-cell patch clamp experiments indicated that cariprazine reduces intrinsic excitability and increases firing irregularity in a cell-type-dependent manner, with regular- and stuttering-type neurons showing the largest effects and delayed firing type neurons being less sensitive. Our prior patch clamp observations already showed that regular firing neurons express mostly outward rectifying K-currents and I_h_, while delayed firing type neurons have strong inward rectifying K_ir_-currents (24). Together, these data support the interpretation that cariprazine primarily modulates intrinsic membrane properties rather than disrupting synaptic coupling within hippocampal networks.

Our electrophysiological findings, supported by conductance-based modeling, implicate Kv1 channel-associated D-type potassium currents as key effectors of the cariprazine response. In patch clamp recordings, cariprazine increased firing irregularity and the prevalence of stuttering phenotypes without major changes in passive membrane parameters. *In silico*, increasing D-current conductance or shifting its activation properties reproduced the reduced excitability and stuttering-like responses, with a stronger impact in the regular-firing model than in the delayed-firing model. These results indicate that modulation of a fast-activating, slowly inactivating potassium current is sufficient to account for the main acute cariprazine effect.

At the molecular level, chronic cariprazine exposure did not alter *Kcna2*/*Kcna3* transcript levels, arguing against transcriptional regulation as the primary driver of the observed electrophysiological changes. Instead, ICC coupled to HCA revealed changes in Kv1.2 protein distribution, including increased Kv1.2-positive puncta number and area in neuronal compartments after cariprazine exposure. This pattern is consistent with post-transcriptional mechanisms such as channel trafficking, clustering, or membrane stabilization, providing a plausible molecular connection to the altered intrinsic excitability (34–36).

The cell-type dependence observed in our recordings may be particularly relevant in the context of hippocampal circuit dysfunction in schizophrenia, where interneuron-mediated coordination of network rhythms is frequently implicated (20, 37). Kv1 channels are prominently expressed in specific interneuron populations and strongly influence spike timing and oscillatory coordination (38). While our culture system does not permit definitive assignment of recorded phenotypes to molecularly defined classes, the preferential sensitivity of neurons with high apparent D-current contribution raises the possibility that cariprazine may disproportionately affect inhibitory elements that shape rhythmic dynamics. Testing this hypothesis in more detailed experimental preparations will be important.

Several limitations should be noted. Cariprazine real-world data were restricted to a relatively short observation window due to its recent clinical introduction; longer follow-up will enable more definitive comparative effectiveness analyses. In addition, the real-world analysis was retrospective and observational, so residual confounding and treatment-selection bias cannot be excluded. The number of first registered suicide-attempt events was also low, limiting power for that endpoint. Sex-and age-stratified analyses were not adequately powered in the present study; future studies should assess whether treatment effects differ by sex and across age groups. Experimentally, primary mouse hippocampal cultures provide mechanistic accessibility but cannot capture the full complexity of human disease biology or chronic *in vivo* drug adaptation. Future work using directly reprogrammed human induced neurons, including patient-derived schizophrenia models, will be instrumental to validate cell-type-specific responses and link drug effects to disease-relevant genetic backgrounds (39–42).

In summary, our study identifies modulation of intrinsic excitability via Kv1/D-type potassium current mechanisms as a candidate cellular pathway through which cariprazine shapes hippocampal neuronal firing while preserving network synchrony. This distinct electrophysiological signature compared with other antipsychotics provides a mechanistic framework to interpret differential clinical profiles and motivates further human-model and circuit-resolved investigations.

## ACKNOWLEDGEMENTS

We thank colleagues at Gedeon Richter Plc. and all members of the HCEMM-SU Neurobiology and Neurodegenerative Diseases Research Group for their valuable comments, discussions, and support during the preparation of this work. We also would like to thank Csaba Kerepesi from SZTAKI, Péter Földi, Tamás Láng, Anna Virág Bakacsi, Vivien Szendi, and Luca Darai for their careful reading of the manuscript and for their valuable feedback and constructive comments.

## AUTHOR CONTRIBUTIONS

M.E.G. and L.D. provided the animals used in the study under the supervision of K.S. A.S., M.E.G., A.A.A., B.K. and K.P. designed and visualized the figures. Á.T., K.L. and A.S. performed and analyzed the computational modeling. M.E.G., A.A.A., B.K. designed and performed the experimental work under the supervision of K.P. M.E.G., A.S. and K.L. designed, supervised and performed patch clamp electrophysiological measurements. M.E.G., S.A. and K.K. performed MEA measurements. I.F. designed the real world statistical analysis and conducted it with the help of A.O. under the supervision of A.B. and C.K. M.E.G., I.F, A.S. and K.P. wrote the manuscript with input from all authors. K.P. and A.S. conceived the study and oversaw overall direction and planning. K.P. supervised the project. All authors provided critical feedback and helped shape the research, analysis and manuscript.

## ETHICS STATEMENT

All experiments involving real-world human data analysis were conducted under the ethical approval number BM/14830-3/2024. Title: Examining the impact of COVID and other diseases of major public health importance on healthcare delivery practices and health outcomes, based on administrative data. Case number: BM/14830-3/2024. Planned duration of the study: 01 July 2024 – 31 July 2027. Issued Budapest, 19 July 2024.

All experiments involving animals described in this study were conducted in accordance with the Hungarian Act of Animal Care and Experimentation (1998. XXVIII), with the directive of the European Parliament and of the Council of 22 September 2010 (2010/63/EU) on the Protection of Animals Used for Scientific Purposes. Protocols were approved by the Animal Care and Use Committee of Eötvös Loránd University and the National Food Chain Safety Office.

## SUPPLEMENTARY INFORMATION

### Supplementary Methods

#### ICD-10 and ATC definitions of baseline characteristics and preselected propensity score covariates

Preselected clinically relevant features for propensity score matching:

In addition to high-dimensional ICD-10 and ATC features selected empirically from the NHIFA database, we included a set of preselected clinically relevant diagnostic and medication covariates in the propensity score models. These variables were selected a priori to improve balance for psychiatric comorbidity, neurological and somatic comorbidities, and baseline concomitant medication use.

For ICD-10 diagnoses, both grouped and individual-code indicators were generated. Grouped features indicated whether a patient had at least one diagnosis within the corresponding ICD-10 code range, while individual-code features represented the occurrence of each separate ICD-10 code within that range. In Supplementary Tables S3–S5, grouped ICD-10 features are denoted as bno_group_ [first ICD code of the group], whereas individual ICD-10 features are denoted as bno_[code].

The preselected ICD-10 diagnostic groups were:

F30–F31, F32–F33, F34–F39, F40–F48, F60–F69, X60–X84, F50–F59, F70–F79, F80–F89, F00–F03/G30–G31, G20–G26, G40–G41, E10–E14, E66, E78, I10–I15, I20–I25, I47–I49, I50, I63–I64, N17–N19, K70–K77, J44, J45, and M00–M99.

For ATC-coded medication use, preselected medication features were generated from two sets of ATC code definitions. First, the following ATC groups were included directly as core medication covariates: N02A, N02B, N02C, M01A, and N06B.

Second, for the following ATC prefixes, all corresponding 5-character ATC codes beginning with the specified prefix were selected and included as medication covariates:

N06A, N03A, N05B, N05C, N04A, A10, C10, B01, and N05A.

In addition, baseline use of selected antipsychotic medications was included as preselected medication covariates using the following ATC codes: olanzapine (N05AH03), quetiapine (N05AH04), paliperidone (N05AX13), risperidone (N05AX08), and clozapine (N05AH02).

Definition of baseline characteristic variables:

Baseline characteristic variables were defined using registered ICD-10 diagnostic codes and ATC-coded prescription records during the baseline/index period. Diagnostic variables were based on the occurrence of the corresponding ICD-10 codes, while medication variables were based on the occurrence of prescriptions with the corresponding ATC codes. Categories were not mutually exclusive; therefore, a patient could contribute to more than one diagnostic or medication category.

ICD-10-defined diagnostic variables:

Substance-use disorder was defined using ICD-10 codes F10–F16 and F18–F19. Smoking or tobacco-use disorder was defined using F17. Depression was defined using F32–F33, bipolar disorder using F30–F31, and anxiety disorder using F40–F43. Organic mental disorder was defined using F00–F09, while dementia was defined using F00–F03 and G30–G31. Epilepsy was defined using G40–G41.

Somatic comorbidities were defined as follows: diabetes using E10–E14, obesity using E66, hypertension using I10–I15, dyslipidemia using E78, ischemic heart disease using I20–I25, heart failure using I50, cerebrovascular disease using I60–I69, and peripheral vascular disease using I70–I74.

Schizophrenia-spectrum diagnostic categories were defined using the following ICD-10 codes: schizophrenia using F20, schizotypal disorder using F21, persistent delusional disorder using F22, acute and transient psychotic disorders using F23, induced delusional disorder using F24, schizoaffective disorder using F25, and other or unspecified schizophrenia-spectrum disorders using F26–F29. Prior registered suicide attempt or intentional self-harm was defined using X60–X84.

ATC-defined medication variables:

Antidepressant use was defined using ATC code N06A. Benzodiazepine use was defined using N05BA, N05CD, and N03AE01. Z-drug use was defined using N05CF. Lithium use was defined using N05AN01, valproate use using N03AG01, carbamazepine use using N03AF01, oxcarbazepine use using N03AF02, and lamotrigine use using N03AX09.

Antiparkinsonian medication use was defined using N04A. Use of selected antipsychotics was defined using the following ATC codes: flupentixol N05AF01, fluphenazine N05AB02, zuclopenthixol N05AF05, olanzapine N05AH03, quetiapine N05AH04, paliperidone N05AX13, risperidone N05AX08, and clozapine N05AH02.

### Supplementary Figures

**Supplementary Figure 1.**
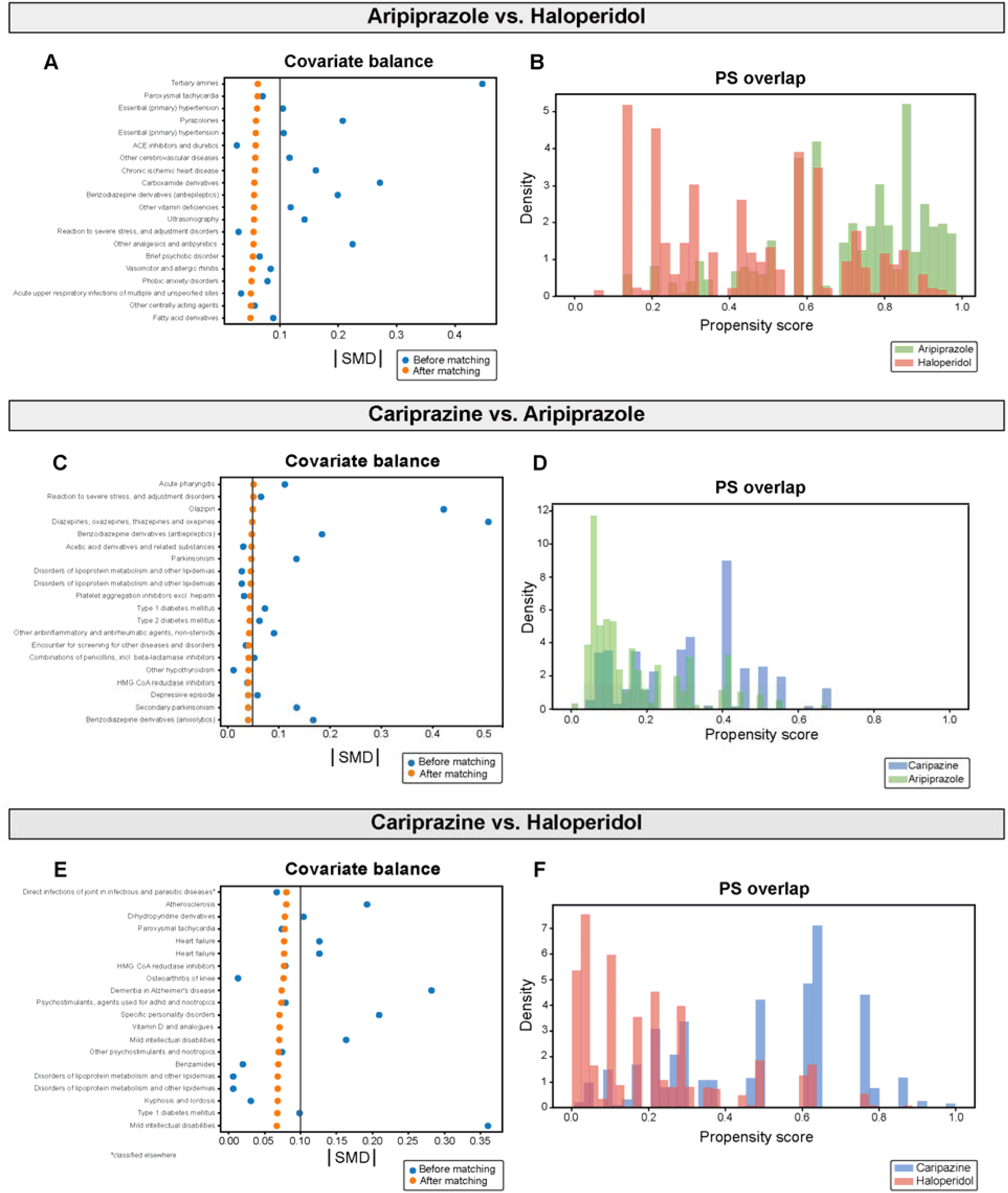
Propensity score matching diagnostics for real-world antipsychotic comparisons. Related to Figure 1. (A, C, E) Covariate balance plots showing the absolute standardized mean differences (|SMD|) of the most imbalanced baseline covariates included in the propensity score model before matching (blue) and after matching (orange) for aripiprazole versus haloperidol (A), cariprazine versus aripiprazole (C), and cariprazine versus haloperidol (E). (B, D, F) Propensity score (PS) overlap plots for aripiprazole versus haloperidol (B), cariprazine versus aripiprazole (D), and cariprazine versus haloperidol (F). Matching reduced covariate imbalance among model-included features and improved comparability between the treatment groups used for the clinical outcome analyses.

**Supplementary Figure 2.**
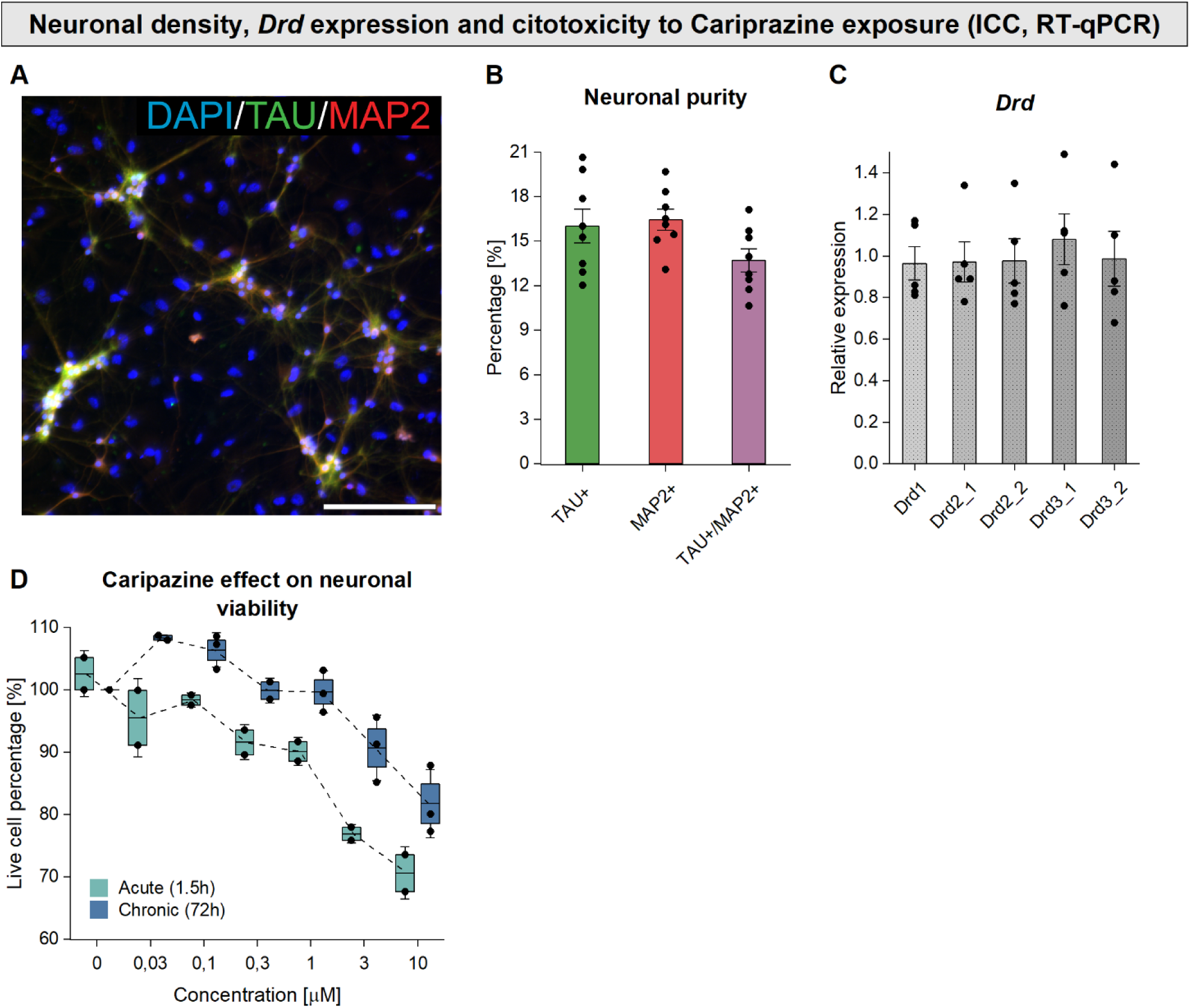
Multimodal characterization of CD1 hippocampal neuronal cultures. Related to Figure 2. (A) Representative high-content microscopy image of hippocampal neuronal cultures (10x magnification, scale bar: 100 µm). Cells were immunostained for the neuronal markers Tau and Map2, with nuclei counterstained using DAPI. (B) Quantification of Tau-positive, Map2-positive, and Tau/Map2 double-positive cells in hippocampal cultures (n = 8). (C) RT-qPCR analysis of dopamine receptor subtype expression (*Drd1* - *Drd3*) in hippocampal neuronal cultures (n = 5). Group differences were evaluated by using Kruskal-Wallis and Dunn’s post-hoc tests. (D) Neuronal viability after acute and chronic CAR treatment with increasing concentrations (acute: n = 14 culture, 2 wells/concentration; chronic: n = 17 culture, 3 well/concentration).

**Supplementary Figure 3.**
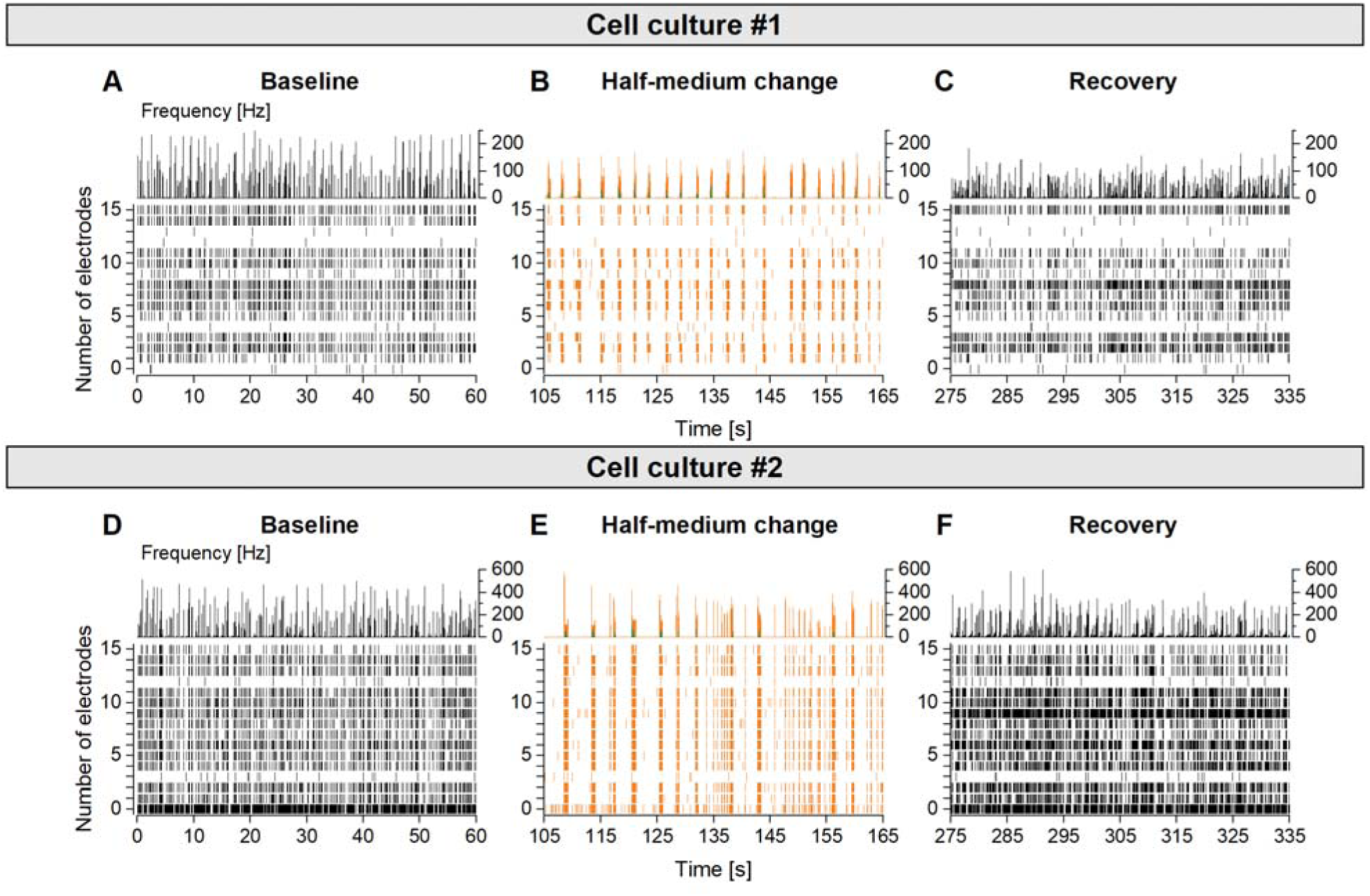
Representative MEA recordings showing transient network perturbation after half-media exchange. Related to Figure 2. (A and D) MEA recordings of baseline spontaneous network activity in independent hippocampal neuronal cultures. Neurons exhibited strong, synchronized network activity characterized by multiunit bursting with variable burst durations. (B and E) Effect of half-media exchange on network activity in the same cultures. Media change transiently reduced overall firing rates, shortened synchronized superburst episodes, and increased burst cycle periods, followed by gradual recovery of activity during the recording period. (C and F) Recovery phase after media exchange, showing normalization of burst duration and frequency toward baseline network activity levels.

**Supplementary Figure 4.**
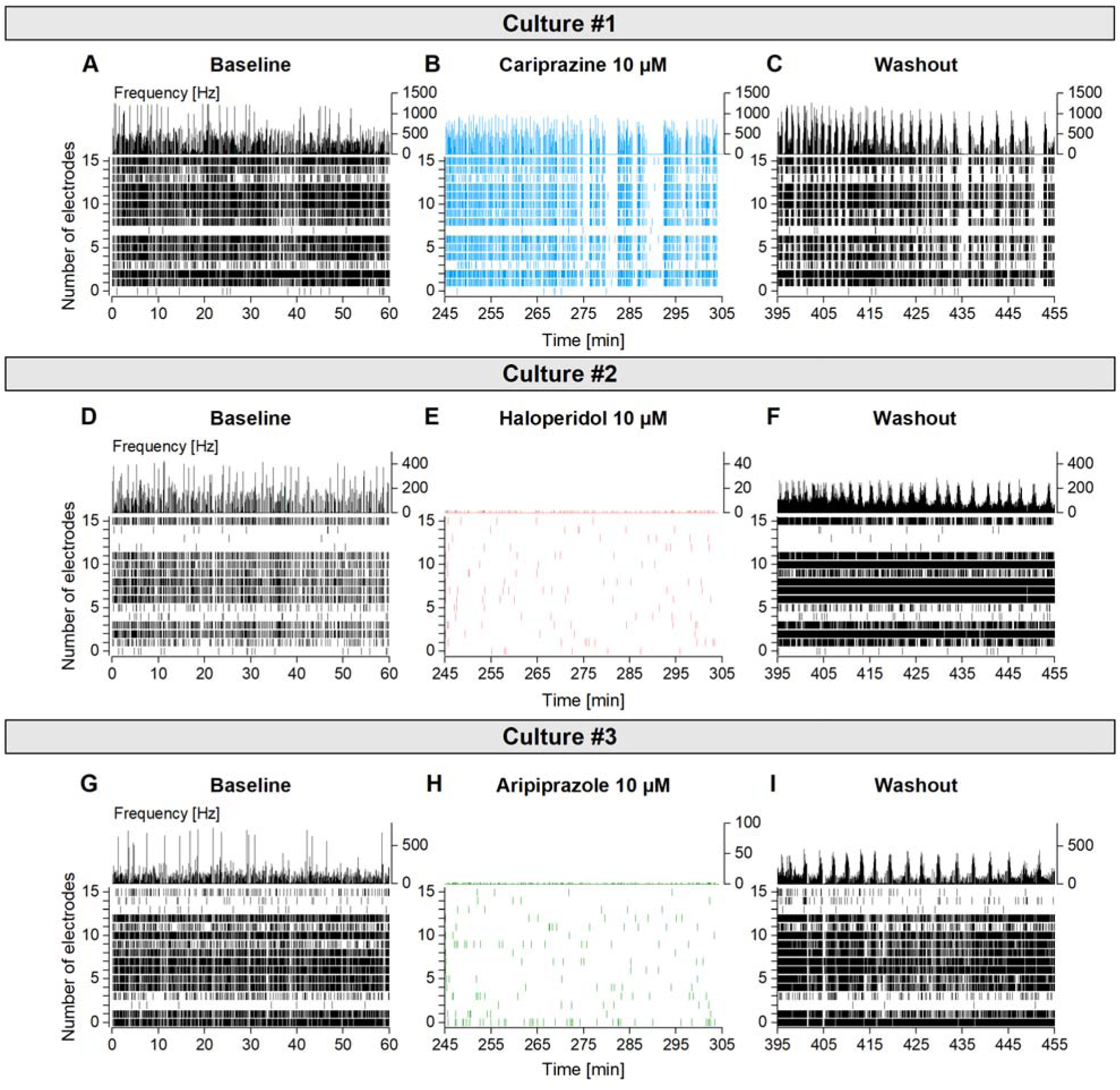
Recovery of neuronal activity after drug application and washout with culture medium. Related to Figure 3. (A, D, G) Hippocampal baseline network activity of three different cultures recorded at the same time using the Axion Maestro Edge MEA system. Firing activity of neurons were monitored for a 1-hour recording period in conditioned culture medium. (B, E, H) Acute antipsychotic action of cariprazine, haloperidol and aripiprazole (10 µM) on network activity in the same three cultures. Cariprazine application (B) decreased the firing rate of hippocampal neurons, but network superbursts were still visible for 1-hour after drug application. Haloperidol and aripiprazole (E, H) nearly eliminated all network activity and synchronous network bursts (superbursts) in the cultures. (C, F, I) Washout of antipsychotics with fresh culture medium. Network activity of hippocampal neurons was restored by the removal of cariprazine, haloperidol and aripiprazole with a decreasing burst frequency by the end of the 1-hour recording session.

**Supplementary Figure 5.**
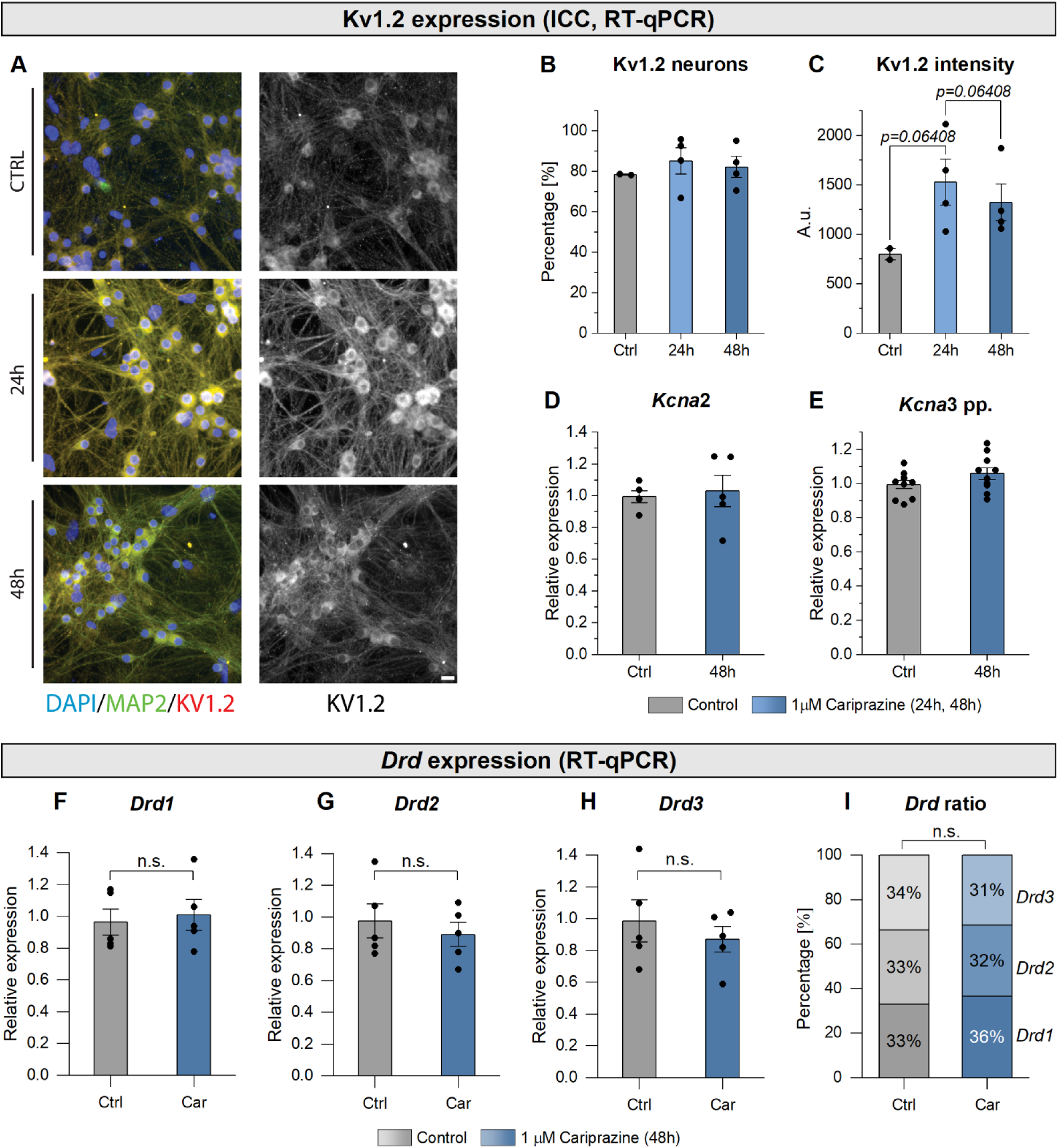
Immunocytochemical detection of Kv1.2 channels and expression of *Kcna* and dopamine receptor (*Drd*) subtype mRNAs following chronic 1 µM Cariprazine treatment. (A) Representative images of hippocampal cultures stained with nuclear (DAPI), neuronal Map2 marker and Kv1.2 channel antibody (20x, enlarged images, scale bar: 12.5 µm) after 24 and 48- hour chronic cariprazine treatment. (B) Percentage of Kv1.2 positive neurons in control (Ctrl) and cariprazine-treated (24h, 48h) conditions. Only responder (cells with higher signal than no-primary antibody control) neurons were counted. (C) Intensity of Kv1.2 staining in control and cariprazine-treated cultures. Cell culture number used for Kv1.2 staining: n = 2 (control), n = 2 (no-primary antibody control), n = 4 - 4 for each condition (24h CAR, 48h CAR). (D and E) Kcna gene expression (mRNA) in control (n = 5) and 1µM cariprazine-treated wells for 48-hours (n = 5) measured by RT-qPCR. (F-H) Dopamine receptor expression measured by RT-qPCR in control (Ctrl) and cariprazine-treated (1µM Car) cell cultures. (I) Ratio of dopamine receptor subtypes in control and cariprazine-treated cell cultures. Group differences were evaluated by using Welch’s two sample t-tests. Dots represent normalized expression values for individual cell cultures. n = 5 for each group (Ctrl, Car). *Drd* subtypes were measured in the same cultures.

### Supplementary Tables

**Supplementary Table 1.**
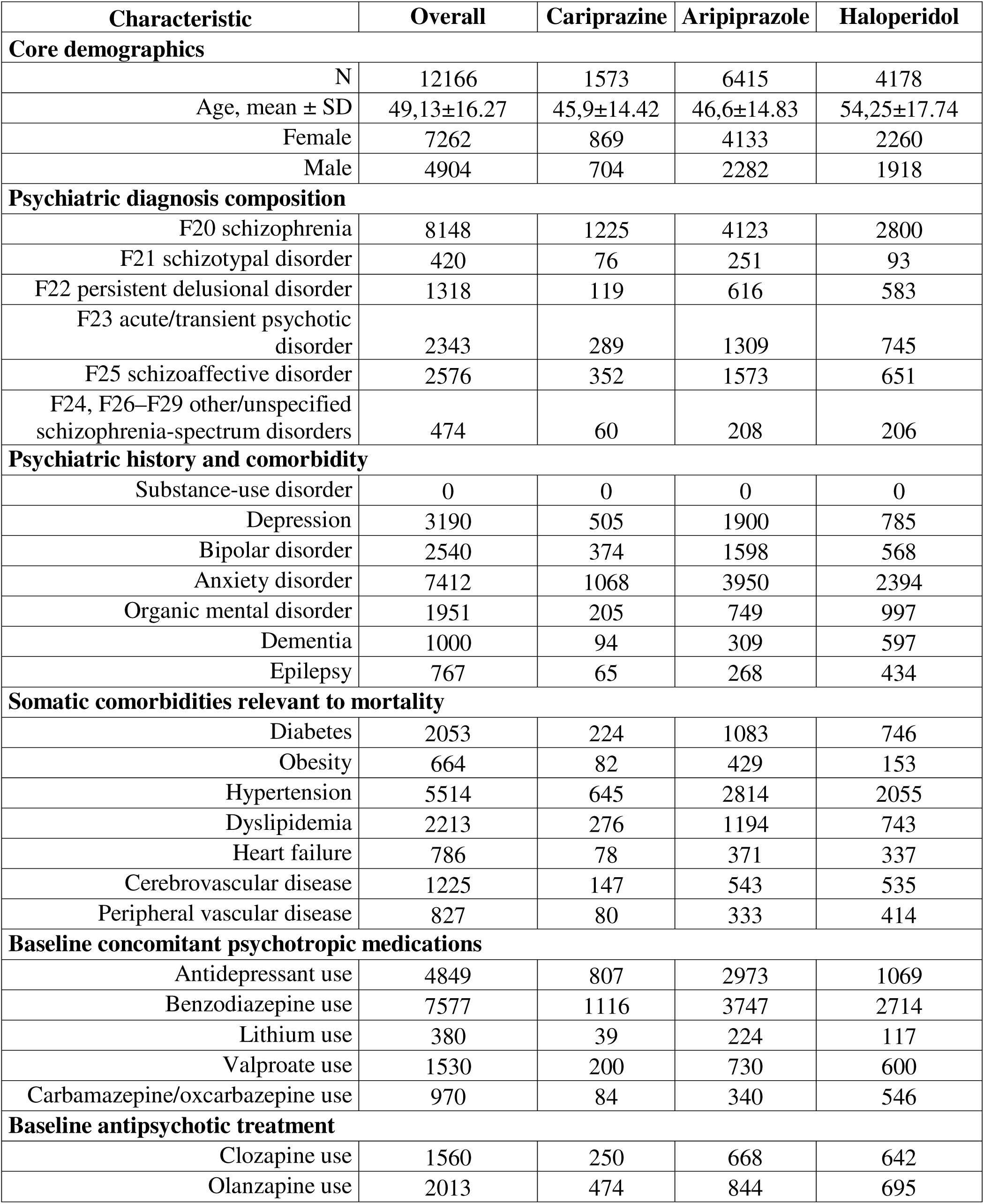
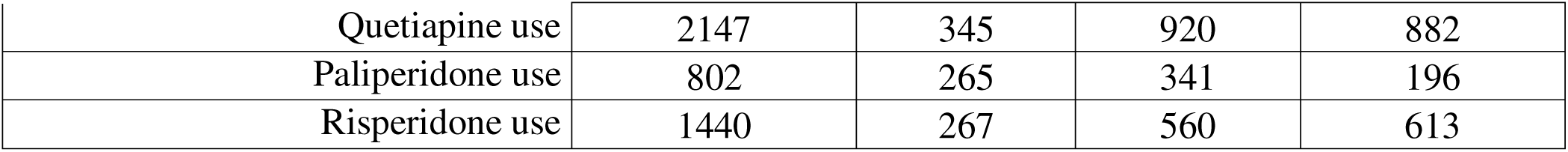
Baseline characteristics of the unmatched nationwide schizophrenia-spectrum cohort. The table shows demographic characteristics, recorded schizophrenia-spectrum diagnoses, psychiatric and somatic comorbidities, and baseline concomitant medication use among patients assigned to cariprazine, aripiprazole, or haloperidol during the index period (from 1^st^ January 2018 to 31^st^ December 2018). Values are presented as counts unless otherwise indicated. Age is reported as mean ± standard deviation. Diagnostic and comorbidity categories were defined using registered ICD codes, and medication variables were defined using prescribed ATC codes during the index period. The full definitions of the comorbidity and medication covariates are provided in Supplementary Methods. Additional baseline characteristics are provided in Supplementary Table S6. Diagnostic categories and comorbidity variables were not mutually exclusive; therefore, counts may sum to more than the total number of patients within a treatment group.

**Supplementary Table 2.** Post-matching covariate balance after pairwise high-dimensional propensity score matching. Balance diagnostics are shown for each pairwise matched comparison between cariprazine, aripiprazole, and haloperidol. Matching was performed separately for each comparison using a 1:1 ratio without replacement and was constrained by sex and 5-year age strata. Median absolute standardized mean differences are reported for variables included in the propensity score model. Additional clinically relevant baseline characteristics reported descriptively in Supplementary Table 1, including variables not fully represented in the final propensity score feature set, were assessed separately for residual imbalance. Residual imbalance was defined as an absolute standardized mean difference greater than 0.1. The sign of the SMD indicates the direction of imbalance according to the order of treatment groups in the matched comparison name. A positive SMD indicates that the covariate was more frequent, or had a higher mean value, in the first-listed drug group, which was considered the treated group. A negative SMD indicates that the covariate was more frequent, or had a higher mean value, in the second-listed drug group, which was considered the comparator group (e.g. in the cariprazine–aripiprazole matched cohort, cariprazine was the treated group and aripiprazole was the comparator group). Baseline characteristic features refer to the clinically relevant variables summarized in Supplementary Table 1. Full SMD values for both propensity score model-included features and baseline characteristic features are provided in Supplementary Tables S3–S5.

| Balance metric | Cariprazine–<br>aripiprazole | Cariprazine–<br>haloperidol | Aripiprazole–<br>haloperidol |
| --- | --- | --- | --- |
| Matched pairs | 1,475 | 1,119 | 2,656 |
| Matched ratio | 1:1 | 1:1 | 1:1 |
| Median abs. SMD among<br>PS-model features | 0.018 | 0.032 | 0.027 |
| Median abs. SMD among<br>Baseline characteristic<br>features | 0.022 | 0.048 | 0.040 |
| Max abs. SMD among<br>PS-model features | 0.050 | 0.080 | 0.062 |
| Max abs. SMD among<br>Baseline characteristic<br>features | 0.168 | 0.173 | 0.081 |
| PS-model features with<br>abs. SMD > 0.1 | None | None | None |
| Baseline characteristic<br>features with abs.<br>SMD > 0.1 | Present | Present | None |
| Antidepressant use | — | 0.117 | — |
| F20 schizophrenia | 0.168 | 0.118 | — |
| Lithium use | –0.110 | –0.173 | — |

**Supplementary Table S3. Post-matching standardized mean differences for the cariprazine–haloperidol matched cohort.**

**Supplementary Table S4. Post-matching standardized mean differences for the aripiprazole–haloperidol matched cohort.**

**Supplementary Table S5. Post-matching standardized mean differences for the cariprazine–aripiprazole matched cohort.**

**Supplementary Table S6. Baseline characteristics of the study cohort.**

**Supplementary Table 7.** Parameters used for *in silico* modeling of single-cell responses from biological neurons under whole-cell patch clamp conditions. Related to Figure 5.

| Current | Na |  | H |  | K<sub>d</sub> |  | M |  | D |  | K<sub>ir</sub> |  |
| --- | --- | --- | --- | --- | --- | --- | --- | --- | --- | --- | --- | --- |
| Type | Reg/<br>Stu | Del | Reg/<br>Stu | Del | Reg/<br>Stu | Del | Reg/<br>Stu | Del | Reg/<br>Stu | Del | Reg/<br>Stu | Del |
| **g (nS)** | 50.000 | 50.000 | 1.5 | - | 350 | 400 | 22 | 18 | 250 | 50 | - | 25.0 |
| **E (mV)** | 55 |  | -38 |  | -78 |  | -78 |  | -78 |  | -78 |  |
| **So (%)** | 30 |  | 30 |  | 30 |  | 50 |  | 50 |  | 50 |  |
| **Ax (%)** | 70 |  |  |  | 70 |  | 50 |  | 50 |  |  |  |
| **De (%)** |  |  | 70 |  |  |  |  |  |  |  | 50 |  |
| **p** | 3 |  | 1 |  | 4 |  | 1 |  | 3 |  | 1 |  |
| **V<sub>m,1/2</sub> (mV)** | -23 |  | -73 |  | -19 |  | -23 |  | -36 |  | -86 |  |
| **V<sub>m,sl</sub> (mV)** | 13 |  | -16 |  | 19 |  | 19 |  | 15 |  | -15 |  |
| **V<sub>h,1/2</sub> (mV)** | -62 |  |  |  |  |  |  |  | -67 |  |  |  |
| **V<sub>h,sl</sub> (mV)** | -16 |  |  |  |  |  |  |  | -10 |  |  |  |
| **τ<sub>m,max</sub> (ms)** | 0.5 |  | 200 |  | 6 |  | 200 |  | 10 |  | 70 |  |
| **τ<sub>m,min</sub> (ms)** | 0.1 |  | 15.0 |  | 0.7 |  | 80.0 |  | 4.0 |  | 8.0 |  |
| **V<sub>tm,1/2</sub> (mV)** | -80 |  | -62 |  | -70 |  | -40 |  | -80 |  | -40 |  |
| **V<sub>tm,sl</sub> (mV)** | 40 |  | 30 |  | 30 |  | 70 |  | 70 |  | 40 |  |
| **τ<sub>h,max</sub> (ms)** | 2 |  |  |  |  |  |  |  | 300 |  |  |  |
| **τ<sub>h,min</sub> (ms)** | 0.5 |  |  |  |  |  |  |  | 200 |  |  |  |
| **V<sub>th,1/2</sub> (mV)** | -86 |  |  |  |  |  |  |  | -90 |  |  |  |
| **V<sub>th,sl</sub> (mV)** | 40 |  |  |  |  |  |  |  | 70 |  |  |  |

